# Alternative splicing of synaptotagmin 7 regulates oligomerization and short-term synaptic plasticity

**DOI:** 10.1101/2025.10.27.684894

**Authors:** Nikunj Mehta, Devin T. Larson, Mitch Wozney, Shweta Mishra, Smrithika Subramani, Simi Kaur, Avani Jain, Edwin R. Chapman

## Abstract

Synaptic plasticity is crucial for learning and memory. The presynaptic calcium sensor synaptotagmin 7 (syt7) regulates aspects of short-term plasticity (STP), but the underlying mechanisms remain unclear. Here, we show that alternative splicing of the syt7 juxtamembrane linker acts as a molecular switch at both biochemical and functional levels. The α and β variants undergo liquid-liquid phase separation to form condensates, while the γ variant forms aggregates. Using iGluSnFR imaging, we found that when expressed at equal levels, these three isoforms also diverge regarding their abilities to regulate two key aspects of STP: paired-pulse facilitation and synaptic depression. STED microscopy showed that all three isoforms form active zone-associated clusters that colocalize with syt1, while MINFLUX super-resolution microscopy resolved syt7 clusters within the active zone, well-positioned to directly control synaptic vesicle dynamics. Thus, alternative splicing might fine-tune STP by differentially impacting syt7 oligomerization.

**Significance Statement:** Neurons communicate via synapses that are often able to dynamically adjust their strength and are hence ‘plastic’. We investigated synaptotagmin 7 (syt7), a key protein that regulates aspects of this synaptic plasticity. We found that alternative splicing functions as a master switch, producing syt7 isoforms that either form liquid condensates or solid aggregates. This biophysical divergence enables the specific fine-tuning of synaptic responses, uncoupling the function of syt7 in two different aspects of plasticity: facilitation and depression. Using STED microscopy, we found that all three syt7 isoforms form active zone-associated clusters that colocalize with syt1, while MINFLUX localized syt7 to regions positioned to modulate transmission. This work highlights a critical link between alternative splicing, protein phase separation, and brain function.

## Introduction

Intricate communication between neurons, via synaptic transmission, forms the foundation of neural circuit function and ultimately underlies cognition and behavior. Many kinds of synapses in the central nervous system are endowed with the ability to rapidly and reversibly change the strength of transmission over timescales ranging from milliseconds to seconds, collectively referred to as short-term plasticity (STP). Synaptic facilitation and depression are two forms of STP, where successive action potentials (AP) lead to a transient increase or decrease in the amplitude of synaptic response, respectively^1–3^. These processes are predominantly governed by presynaptic mechanisms that modulate the efficacy of synaptic vesicle (SV) fusion with the plasma membrane^4–7^.

SV docking and fusion are regulated by members of the synaptotagmin (syt) family of proteins^8^, characterized by the presence of tandem C2 domains (C2A and C2B) that often serve as Ca^2+^-binding modules^9–11^, a juxtamembrane (jxm) linker, and a single transmembrane domain (TMD) with a short N-terminal segment (Fig. 1A). The most studied isoform, syt1, is localized to SVs^12^ where it plays a role in: SV docking^7,13–15^, clamping release under resting conditions^16–18^, and triggering rapid, synchronous exocytosis in response to increases in intracellular calcium concentration ([Ca^2+^]_i_) following an AP^16–20^. Another member of this family, syt7, has a similar domain structure but undergoes proteolytic processing to liberate the cytoplasmic domain^21^, which is then anchored to the presynaptic plasma membrane via palmitoylation^22^.

**Figure 1.**
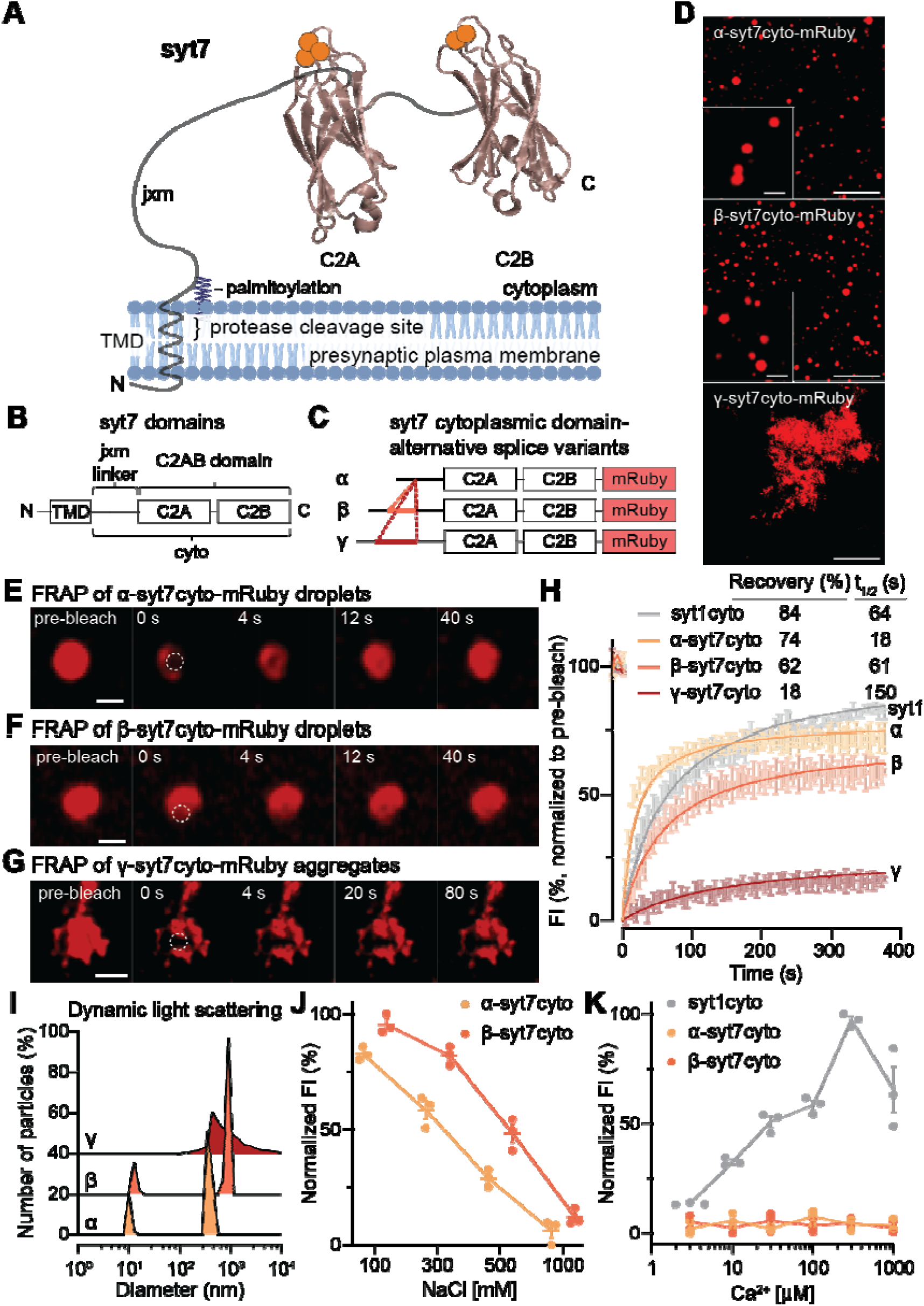
Alternative splicing regulates the oligomerization of synaptotagmin 7. (**A, B**) Illustration of synaptotagmin 7 (syt7), indicating: γ-secretase cleavage within the transmembrane domain (TMD), palmitoylation of the juxtamembrane linker (jxm) that undergoes alternative splicing, and tandem C2 domains, C2A and C2B, that bind Ca^2+^. The membrane was drawn using BioRender, the structures for C2A and C2B (PDB ID: 6ANK) were rendered using UCSF Chimera; Ca^2+^ ions are shown as orange spheres. (**C**) Schematic diagram of the cytoplasmic domain of syt7 (syt7cyto) tagged with a C-terminal mRuby, along with the alternative splice variants: α, β, and γ, which differ only in their jxm linkers. (**D**) α- and β-syt7cyto-mRuby form droplets, whereas γ-syt7cyto-mRuby forms aggregates under the same buffer conditions ([protein]: 10 µM α-, β-; 3 µM γ-syt7cyto-mRuby). Zoomed-in α- and β-syt7cyto-mRuby droplets are shown as insets to emphasize the circular shape of droplets. Scale bar, 10 µm. Inset scale bar, 2 µm. (**E-G**) Time series of fluorescence recovery after photobleaching (FRAP) of (E, F) α- and β-syt7cyto-mRuby droplets and (G) γ-syt7cyto-mRuby aggregates. Dotted circle shows the area, with the highest intensity of photobleaching, within the droplet and aggregate. Scale bars: (E, F), 1 µm; (G), 2.5 µm. (**H**) Quantification of fluorescence intensity (FI) from the FRAP time series for the α-, β-, and γ-syt7cyto-mRuby splice variants shown in yellow, orange, and red, respectively. For comparison, FRAP data of jxm-syt1cyto-GFP are replotted, from our previous work^29^, in grey. Data were fitted using a hyperbolic function, and the calculated recovery (%) and t_1/2_ (s) values are shown in the inset. (**I**) Number of particles (relative%), as a function of diameter (nm), of syt7cyto splice variant droplets/aggregates determined via dynamic light scattering (DLS). (**J**) Normalized FI of α- and β-syt7cyto-mRuby as a function of [NaCl]. (**K**) Same as (J), but as a function of [Ca^2+^]. For comparison, [Ca^2+^] titration of jxm-syt1cyto-GFP was replotted, from ref. ^29^, in grey. Three independent trials with 4-10 FOVs per condition were analyzed, data are plotted as mean ± SEM. The buffer used in (D-J) is 25 mM Tris-HCl (pH 7.4), 100 mM NaCl, and 3% PEG 8000; the same buffer, but lacking PEG 8000, was used in panel (K).

Syt7 has emerged as a crucial protein essential for two forms of STP: it is required for paired-pulse facilitation (PPF), and it endows synapses with the ability to resist synaptic depression during stimulus trains^5,6,21^. Syt7 senses residual [Ca^2+^]_i_ ^3^, which lingers after an AP, and mediates PPF, owing to its significantly higher Ca^2+^ affinity and slower kinetics, as compared to syt1^23,24^. In addition to playing a key role in STP, syt7 also supports asynchronous release (neurotransmitter release tens to hundreds of milliseconds after an AP) during train stimulus^21,25–27^. More recently, syt7 was proposed to mediate PPF and resistance to synaptic depression by regulating the activity-dependent docking of SVs^25^. In short, while syt1 provides the synapse with speed and fidelity, syt7 endows synapses with adaptability and endurance.

An emergent area of interest is the potential for syts to form oligomers^28^ and whether oligomerization impacts their function. Indeed, syt1 oligomerizes via its jxm linker to undergo liquid-liquid phase separation (LLPS)^29^, and this biochemical property strongly influences both synchronous and spontaneous synaptic transmission^30^. Whether syt7 also undergoes LLPS, and how self-association relates to its specific roles in asynchronous release and STP remain largely unexplored. There is some evidence that syt7 can multimerize, but this was proposed to be mediated via its tandem C2-domains^31^. Interestingly, the jxm linker of syt7 also contains an intrinsically disordered region (IDR) that is potentially conducive to phase separation. Strikingly, this jxm linker is subject to extensive alternative splicing, and three major variants - α, β, and γ - have been described^32^. However, biochemical and functional differences between these splice variants have yet to be investigated. Finally, while the localization of syt1 to SVs is well established^9^, the precise localization of syt7 in the plasma membrane of axons is unknown; addressing its spatial organization is needed to understand how syt7 regulates the SV cycle.

The current study aims to investigate the ability of syt7 to oligomerize and to address whether alternative splicing regulates self-association and aspects of STP. By employing iGluSnFR imaging^33^, we directly measured glutamate release from cultured neurons and assessed the impact of three syt7 splice variants: α, β, and γ, on STP. Furthermore, we utilized stimulated emission depletion (STED) imaging of all three splice variants and two-color three-dimensional minimal photon flux (MINFLUX) super-resolution imaging^34,35^ to probe the nanoscale organization of syt7 in presynaptic nerve terminals. We report that two splice variants undergo LLPS while one variant forms aggregates. Importantly, these variants also uncoupled the function of syt7 in PPF versus synaptic depression, revealing a functional switch that is subordinate to alternative splicing. Finally, STED microscopy showed that α-, β-, and γ-syt7 all form clusters that colocalize with syt1 at bassoon-positive active zones; MINFLUX imaging further resolved the nanoscale positioning of α-syt7 clusters and confirmed that they are positioned to regulate SV dynamics.

## Results

### Alternative splicing regulates syt7 self-association

Syt7 undergoes alternative splicing in the region that encodes the jxm linker, which connects its calcium-sensing tandem C2-domains to the presynaptic plasma membrane via palmitoylation (Fig. 1A, B; Supplementary Table 1). It is currently thought that there are three major splice variants of syt7-α, β, and γ (Fig. 1C)^32,36^. The expression of these isoforms is developmentally and regionally regulated^36^, and it has been proposed that they might function differentially during insulin secretion from pancreatic β cells^37^. The discovery that syt1 undergoes LLPS via its jxm linker^29^, and that this interaction is crucial for function^30^, prompted us to explore whether syt7 also undergoes LLPS. As noted above, the syt7 linker has an IDR and, again, its length and composition are impacted by alternative splicing (Supplementary Table 1). Using mRuby as a C-terminally tagged fluorescent label on the cytosolic (cyto) domain of syt7 (Fig. 1C), we found that the α and β variants formed droplets in aqueous buffer, whereas the γ variant formed amorphous and disordered aggregates under the same conditions (Fig. 1D). To further validate the differential homo-oligomerization of syt7 splice variants, we performed fluorescence recovery after photobleaching (FRAP) on the α- and β-syt7cyto droplets, and γ-syt7cyto aggregates (Fig. 1E-H). FRAP data were fitted to hyperbolic functions to calculate the extent and t_1/2_ of recovery. After bleaching within protein droplets and aggregates (bleaching diameter between 0.8-2 µm), α- and β-syt7cyto droplets rapidly recovered (t_1/2_ = 18, and 61 s, respectively; yellow and orange traces; Fig. 1E,F,H), whereas γ-syt7cyto aggregates recovered slowly (t_1/2_ = 150 s; red trace; Fig. 1G,H). The extent of recovery of γ-syt7cyto aggregates was extremely low (18% recovery), as compared to α- and β-syt7cyto droplets (74 and 62% recovery, respectively; Fig. 1E-H), supporting the idea that α- and β-syt7cyto undergo LLPS to form protein droplets, while γ-syt7cyto again tends to form aggregates. For comparison, FRAP of syt1cyto (80-421 residues), from our previous study^29^, is replotted in grey, and exhibited 84% recovery and a t_1/2_ of 64 s (Fig. 1H). To assess the effect of molecular crowding on syt7 splice variants, we characterized droplet formation by each syt7 variant as a function of increasing [PEG 8000] and [protein]. Phase diagrams in Supplementary Fig. 1A-C indicate that β-syt7cyto has a slightly higher propensity to form droplets than α-syt7cyto (at 3 µM [protein] and 3% PEG 8000, and 10 µM [protein] and 1% PEG 8000), whereas γ-syt7cyto forms aggregates under all [protein] and [PEG 8000] conditions. As a control, mRuby and syt7C2AB-mRuby, lacking the jxm linker, did not form droplets under the same conditions (Supplementary Fig. 1D-F), further indicating that the jxm linker drives self-association. Additionally, α-, as well as β-syt7cyto droplets, undergo homotypic fusion, another characteristic property of LLPS droplets (Supplementary Fig. 1G, H).

A classic signature of protein aggregation is high hydrophobicity. Notably, sequence analysis of the jxm linker of the γ-syt7 splice variant showed a higher hydropathicity index (a high index indicates high hydrophobicity) as compared to the jxm linkers of α- and β-syt7 (Supplementary Fig. 2A-C). Size characterization of protein droplets and aggregates by dynamic light scattering (DLS) indicated that the α- and β-syt7cyto droplets had two peaks corresponding to monomeric and higher-order structures, whereas γ-syt7cyto aggregates had a wide range of large diameter structures (Fig. 1I) (note: the DLS instrument cannot distinguish particle sizes >1 µm). As controls, mRuby and syt7C2AB-mRuby showed single monomeric peaks (Supplementary Fig. 3A), establishing that neither the fluorescent tag nor the two C2-domains undergo LLPS under these conditions. To identify the types of interactions within α- and β-syt7cyto droplets, we challenged the droplets with increasing ionic strength and observed that they progressively dissolved (Fig. 1J). These results suggest that electrostatic interactions contribute to their phase separation. In sharp contrast, γ-syt7cyto aggregates were not affected by increasing ionic strength (Supplementary Fig. 3B). We previously found that Ca^2+^ promotes syt1 droplet formation, so we examined the effects of Ca^2+^ on syt7 droplets. As shown in Fig. 1K and Supplementary Fig. 3C, Ca^2+^ did not promote oligomerization of any of the three splice variants. The effect of Ca^2+^ to promote syt1 droplet formation is included for comparison (grey trace; data are replotted from ref. ^29^). Collectively, these data show that the β variant has a somewhat greater propensity to undergo LLPS versus α, while the γ variant forms more rigid aggregates.

### Alternative splicing and subcellular localization determine the extent and kinetics of recovery of syt7 after photobleaching

To examine syt7 LLPS in cells, we expressed the cytoplasmic domains of each of the three mRuby-tagged variants in hippocampal neurons and challenged them with 10% 1,6-hexanediol (1,6-HD), an aliphatic alcohol that disrupts weak hydrophobic interactions within droplets to dissolve them. As shown in Supplementary Fig. 4A,B, α- and β-syt7cyto formed droplets in the soma and neurites. Upon addition of 1,6-HD, the condensates dissolved, in agreement with our biochemical observations that α- and β-syt7cyto undergo LLPS. In contrast, 1,6-HD had no effect on clusters formed by γ-syt7cyto, consistent with the idea that this variant forms aggregates in cells (Supplementary Fig. 4C). These observations lead to the question of LLPS in the context of membrane-associated proteins; in this case, we reiterate that after processing, syt7 is attached to the plasma membrane via palmitoylation^21,22^. To address two-dimensional (2D) LLPS on a membrane, we attached His-tagged α-syt7cyto-mRuby droplets to the surface of giant unilamellar vesicles (GUVs) using an 18:1 DGS-NTA (Ni) lipid. Supplementary Fig. 5A,B shows that α-syt7cyto-mRuby assembled into LLPS patches on a GUV, to form a ‘phase-separated surface’.

We next characterized the full-length (fl) versions of each syt7 splice variant. Each variant was fused to a C-terminal HaloTag (labeled with JF549) and overexpressed in HEK293T cells (Fig. 2A, Supplementary Fig. 6A,B) and rat hippocampal neurons (Fig. 2E,F, Supplementary Fig. 7A,B). In both cases, the fusion proteins were largely targeted to the plasma membrane, where they were subjected to FRAP experiments. In HEK cells, α- and β-syt7-fl recovered 71% and 69%, respectively (Fig. 2B,C), and the t_1/2_ values are provided in Supplementary Table 2. In the presence of 10% 1,6-HD, recovery of both α- and β-syt7-fl was <26% (Fig. 2A-C, Supplementary Fig. 6A), indicating that these proteins were unable to reassemble into LLPS-mediated patches. Analogous to the *in vitro* FRAP experiments of γ-syt7cyto aggregates in Fig. 1G,H, in cells, γ-syt7-fl recovered only 21% of the signal under control conditions, while no significant changes were observed with 1,6-HD treatment (Fig. 2D, fig S6B; % recovery = 15, t_1/2_ values are provided in Supplementary Table 2), consistent with the conclusion that this variant does not form liquid-like dynamic assemblies.

**Figure 2.**
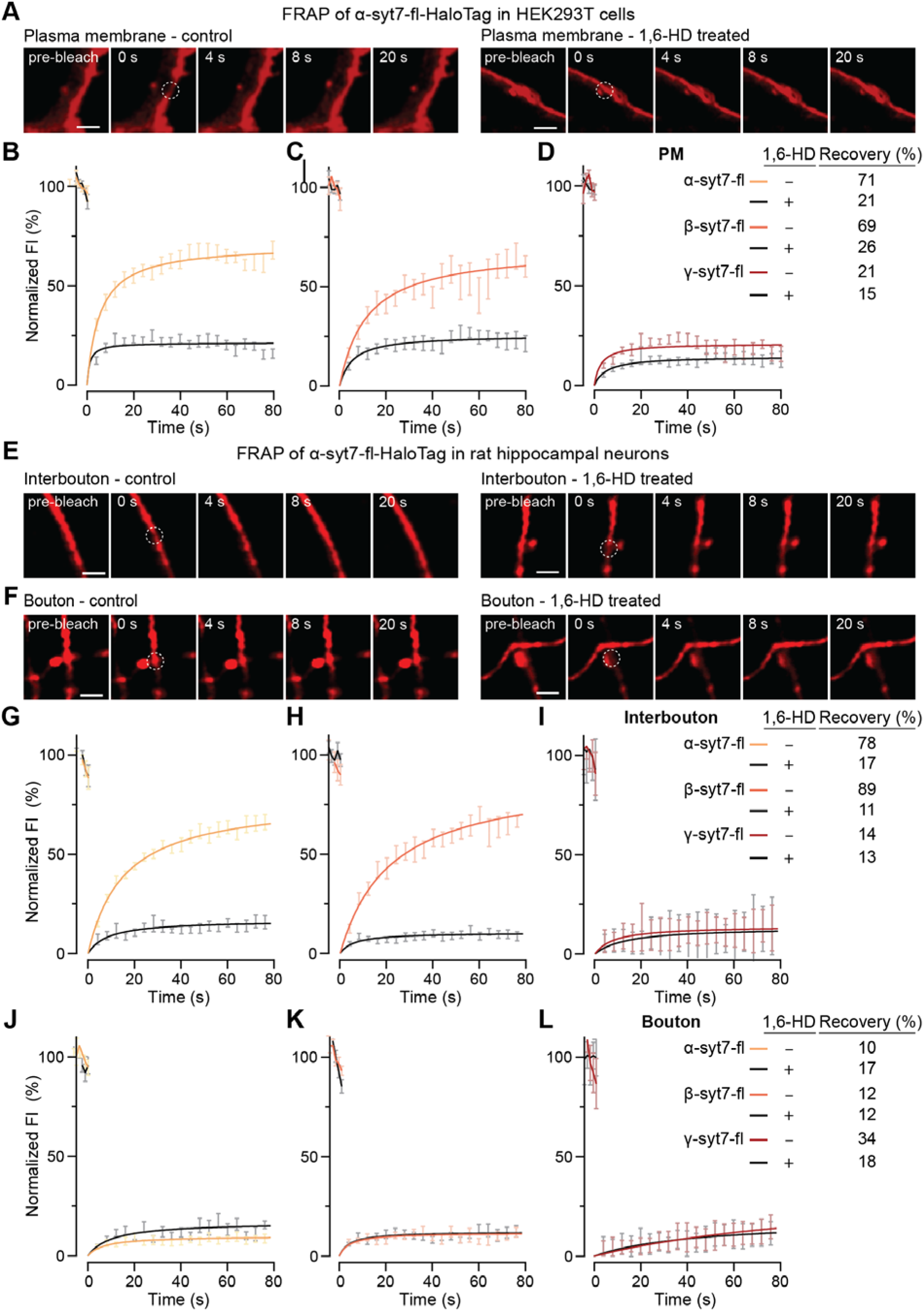
Extent and kinetics of recovery from photobleaching depend on alternate splicing and subcellular localization of syt7. (**A**) Time series of FRAP at the plasma membrane of HEK293T cells expressing α-syt7-full-length (fl)-HaloTag, under control and 10% 1,6-hexanediol (1,6-HD; an aliphatic alcohol that disrupts phase separation of proteins) conditions, respectively. Scale bar, 2 µm. (**B-D**) Quantification of FRAP in HEK293T cells expressing α-, β-, or γ-syt7-fl-HaloTag splice variants in yellow, orange, and red, respectively; the black traces in each panel indicate FRAP experiments with 10% 1,6-HD treatment. Data were fitted using a hyperbolic function to calculate recovery (%) and t_1/2_ (s), as shown in the inset and Supplementary Table 2, respectively. (**E,F**) FRAP time series of α-syt7-fl-HaloTag along axons (interbouton) and within boutons, respectively, in rat hippocampal neurons, under control and 10% 1,6-HD conditions. Scale bar, 2 µm. (**G-I)** and **(J-L**) Same as (B-D), but within interbouton segments versus boutons, respectively. Inset shows %recovery; Supplementary Table 3 indicates calculated t_1/2_ (s) values. N = 15-30 FRAP traces across each condition from three independent trials; data are represented as mean ± SEM. Dotted circles indicate bleached area in FRAP experiments. HaloTag-fusion proteins were labeled with JF549. Representative FRAP images, across different cells and conditions, were brightness- and contrast-adjusted.

Extending these observations to rat hippocampal neurons, the FRAP profiles of α- (Fig. 2E,F), β- (Supplementary Fig. 7A), and γ-syt7-fl (Supplementary Fig. 7B) were similar to HEK cells only within interbouton segments (Fig. 2G-I; % recovery under control and 1,6-HD conditions, respectively, for α: 78, 17, β: 89, 11, and γ: 14, 13; the t_1/2_ values are provided in Supplementary Table 3). These data are consistent with previous observations that syt1 and other presynaptic proteins form clusters on the plasma membrane of neurons^38^. Surprisingly, the extent of recovery was severely hindered in boutons, and no significant changes were observed upon treatment with 1,6-HD (Fig. 2J-L; % recovery under control and 1,6-HD conditions, respectively, for α: 10, 17; β: 12, 12; and γ: 34, 18; the t_1/2_ values, are provided in Supplementary Table 3), revealing limited diffusion within boutons (see Discussion). We also conducted FRAP experiments of the α variant in the soma, dendrites, and the axon initial segment (AIS; Supplementary Fig. 8A-D), and observed limited recovery in all three locations, and little to no effect of 1,6-HD (Supplementary Fig. 8A,B,D). Together, these results demonstrate that alternative splicing and subcellular localization alter the extent and kinetics of recovery of syt7 after photobleaching.

### Alternative splicing of syt7 affects paired-pulse facilitation

To determine whether alternative splicing regulates the function of syt7, we examined presynaptic aspects of PPF and synaptic depression in cultured hippocampal neurons using iGluSnFR, an optical sensor that directly reports glutamate release^33^. After transducing neurons with a lentivirus to express the sensor, we examined PPF by triggering two APs at 20 Hz (50 ms interstimulus interval (ISI)) and calculated the ratio of the second to the first iGluSnFR response (Fig. 3A). As observed previously^5,6,21,25^, cultured WT mouse hippocampal neurons exhibited a paired-pulse ratio (PPR) of 1.24 ± 0.02, whereas syt7KO neurons exhibited paired-pulse depression, with a ratio of 0.63 ± 0.02 (see Fig. 3A,B for raw traces and quantification, respectively). We then expressed each of the three splice variants in a syt7KO background; α-syt7 was expressed at a level that functionally matched WT PPF, while β- and γ-syt7 were matched to the α-syt7 protein levels (Supplementary Fig. 9A-D for Western blots). Interestingly, all three variants, when expressed at equal levels, rescued PPF (Fig. 3A,B), albeit to different degrees and via different mechanisms. The β variant yielded a strong second response, even greater than in WT, with a normal first response, to enhance PPF (ratio: 1.69 ± 0.03; Supplementary Fig. 10A,B, Fig. 3A,B). In contrast, both the α and γ variants suppressed the first response, with γ having a particularly dramatic effect (Supplementary Fig. 10A). This suppression is reduced in the second response, resulting in PPF (ratios, α: 1.27 ± 0.02, γ: 2.42 ± 0.05; Fig. 3A,B, Supplementary Fig. 10B). These findings demonstrate clear functional differences between the three alternatively spliced isoforms of syt7 when expressed at equal levels.

**Figure 3.**
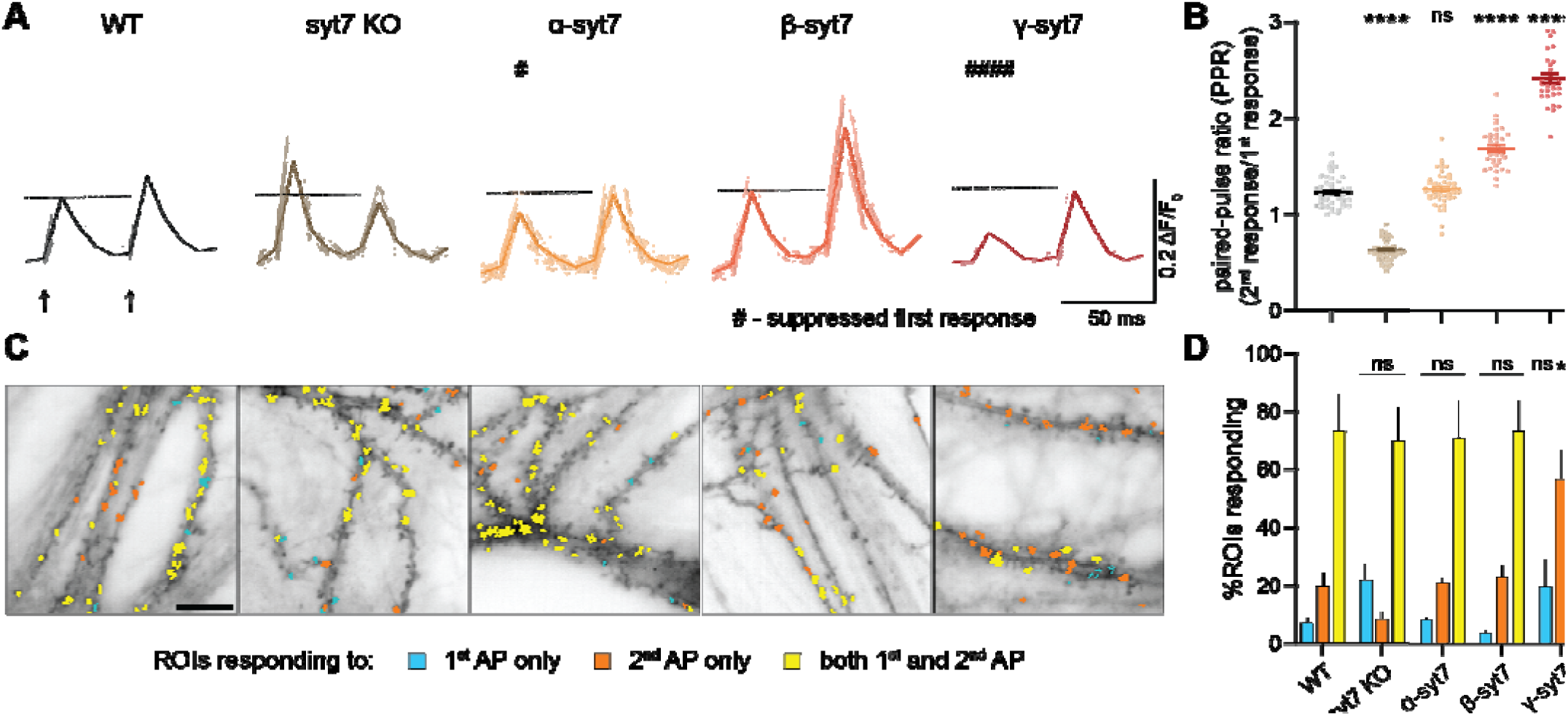
Alternative splicing of syt7 regulates paired-pulse facilitation. (**A**) Representative traces of SF-iGluSnFR.S72A (henceforth referred to as iGluSnFR) responses (ΔF/F_0_) from a field of view (FOV) of mouse hippocampal neurons after two stimuli at 20 Hz (indicated by two arrows); the imaging frequency was 100 Hz. Average traces from a FOV of wild-type (WT), syt7KO, and syt7KO transduced with equal amounts of α-, β-, or γ-syt7fl are in black, brown, yellow, orange, and red, respectively, with lighter shades representing traces from individual regions of interest (ROIs). The dashed line corresponds to the peak of the first response in WT condition. #*P* < 0.05; ####*P* < 0.0001 indicates suppressed first response. (**B**) Quantification of paired-pulse ratios (PPR) of peak iGluSnFR responses for all five conditions tested in (A). Note that the first response of γ-syt7 is strongly suppressed as compared to WT, and hence, PPR appears synthetically high. (**C**) Representative FOV of all five conditions from panel (A) showcasing ROIs that respond to only first (cyan), only second (orange), and both (yellow) APs. Note: the number of ROIs that respond to both the first and second APs are significantly lower in the γ-syt7 condition compared to any other condition tested, owing in part to an increased failure rate during the first AP. Scale bar, 10 µm. (**D**) Quantification of (C), demonstrating % of ROIs responding to only the first, second, or both APs. Note that α-syt7 protein expression was functionally matched to WT levels to elicit a similar PPR, and β-, and γ-syt7 expression levels were kept similar to α-syt7 (Supplementary Fig. 9A-D). Number of FOVs analyzed: 44, 48, 48, 35, and 30 for WT, syt7KO, α-, β-, or γ-syt7 conditions, respectively, across three or more independent culture preparations; each FOV contains 50-200 ROIs; data are represented as mean ± SEM. To determine statistical significance, one-way analysis of variance (ANOVA) with Tukey’s correction for multiple comparisons was used in (B); two-way ANOVA with Tukey’s correction for multiple comparisons in (D). ns, not significant; \**P* < 0.05; \*\*\*\**P* < 0.0001; ^##^*P* < 0.01. Comparisons to WT condition are indicated in (B); comparisons to WT condition for first, second, and both %ROIs responding in (D); full statistics are provided in Data S1.

To determine whether the splice-variant-dependent effects on PPF were maintained across different ISIs, we measured PPR over a range of ISIs from 50 to 1000 ms, corresponding to 20, 10, 5, 2, and 1 Hz stimulation. As reported in Vevea et al. (2021), and Jackman et al. (2016)^6,21^, WT neurons showed facilitation at short ISIs (50 and 100 ms ISI) and this decayed as the intervals between stimuli were increased, whereas syt7KO neurons exhibited paired-pulse depression at all time points (Supplementary Fig. 11A). Expression of each syt7 splice variant restored facilitation at short ISIs, but to different extents, with γ-syt7 producing the largest PPR, followed by β- and then α-syt7. These isoform-specific differences were most apparent at shorter ISIs and diminished at longer intervals. The PPF decay kinetics were similar for WT and the α and β variants; all were well-fitted by a single exponential yield time constants in the 129-212 ms range. However, the γ-syt7 variant differed; this decay was well-fitted by a double exponential function and yielded time constants of 50 and 1280 ms (See Supplementary Table 4 for quantification). These findings indicate that alternative splicing can tune both the magnitude and, for one isoform, the kinetics of syt7-dependent PPF.

To investigate the suppression of the first response by the α and γ variants, we leveraged the spatial readout afforded by iGluSnFR to map the regions of interest (ROIs) responding to only the first (blue), only the second (orange), or both APs (yellow; Fig. 3C). This analysis showed that γ-syt7 is, again, unusual; this variant yielded a significant reduction in the fraction of synapses that responded to both APs, as compared to all the other conditions. In addition, a significant population of synapses were recruited only upon the second AP (Fig. 3C,D), as evidenced by the appearance of iGluSnFR responses that were silent during the first AP. In contrast, neurons expressing α- or β-syt7 displayed a similar number of synapses that responded to both APs.

Delving further, we analyzed the PPR by examining the ROIs that only responded to both APs (Supplementary Fig. 12A). This enabled us to address whether syt7 variants affect PPF by increasing glutamate release, independent of synapse recruitment during the second AP. In WT neurons, as well as KO neurons expressing each of the three syt7 splice variants equally, there was an increase in glutamate release in response to the second stimulus (Supplementary Fig. 12A; note: the number of synapses were the same across splice variants (Supplementary Fig. 13A,B)). These findings reveal that syt7-mediated PPF involves, to some degree, strengthening the output of reliable, active boutons. Future work, marking all synapses, is needed to quantitatively address the contribution of synaptic recruitment.

### Alternative splicing of syt7 impacts synaptic depression

Loss of syt7 accelerates and deepens synaptic depression, due to a reduction in the SV replenishment rate (SVRR)^21,39^. In the next series of experiments, we addressed the roles of the three splice variants in this form of STP. We again turned to iGluSnFR to directly measure presynaptic aspects of depression, but now during train stimulation (50 APs at 20 Hz). Average iGluSnFR traces and amplitudes are shown in Fig. 4A-B, confirming faster and greater depression during the train in syt7KOs as compared to WT^5,6,21,25^ (Supplementary Fig. 14A). Strikingly, in our equal-expression-level paradigm, only the β variant rescued the depression phenotype. Of note, neurons expressing this isoform performed even better than WT, with larger amplitude glutamate signals that resisted depression (Fig. 4A-B). Our analysis reports absolute amplitudes, as normalizing the responses in the train to the first response would obscure biologically important differences when the first response, especially in the case of the γ variant, is strongly and selectively suppressed. Neurons expressing α were largely indistinguishable from the KO; γ was similar to α but had slightly enhanced depression (Fig. 4A-B). We then plotted the cumulative iGluSnFR signal (Fig. 4C) and calculated the SVRR from the slope of a linear curve fitted to the final 30 APs of the train (Fig. 4D). Only β-syt7 was able to rescue SVRR as compared to the WT condition, while α and γ were indistinguishable from KO (Fig. 4D). In summary, under similar expression levels of the three syt7 splice variants in cultured neurons, only the β splice variant rescues both the PPF and depression phenotypes in the KO, while all three variants rescue PPF (albeit via somewhat different mechanisms). Hence, alternative splicing uncouples the roles of syt7 in these two forms of short-term plasticity.

**Figure 4.**
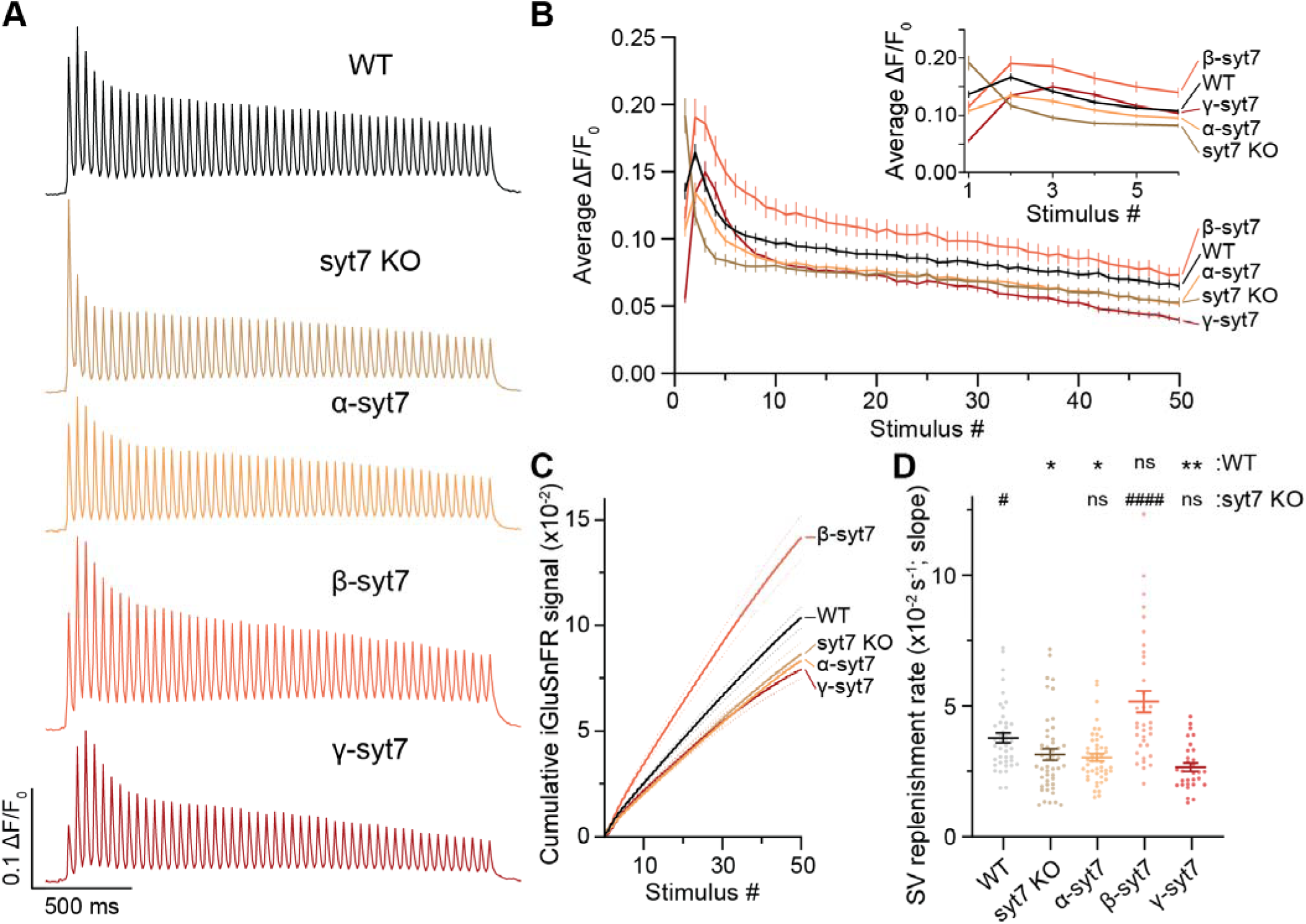
Alternative splicing of syt7 differentially regulates resistance to synaptic depression and synaptic vesicle replenishment rate. (**A**) Average iGluSnFR responses (ΔF/F_0_) from mouse hippocampal neurons during high frequency stimulation (HFS; 50 APs at 20Hz) for WT, syt7KO, and syt7KO that expressed equal levels of α-, β-, or γ-syt7-fl, represented in black, brown, yellow, orange, and red, respectively. (**B**) Average amplitude of iGluSnFR responses (ΔF/F_0_) after HFS in (A). Inset shows first six responses for all conditions. (**C**) Average cumulative iGluSnFR signal for all five conditions plotted versus stimulus number. β-syt7-fl enhances while α- and γ-syt7-fl fail to rescue total glutamate release. (**D**) Synaptic vesicle replenishment rates (SVRR) were calculated from the slope of the linear fit of the last 30 APs of the cumulative release plot (C). Number of FOVs analyzed: 44, 48, 48, 35, and 30 for WT, syt7KO, α-, β-, or γ-syt7 conditions, respectively, across ≥3 culture preparations; each FOV contains 50-200 ROIs; data are mean ± SEM. To determine statistical significance, one-way analysis of variance (ANOVA) with Kruskal-Wallis test with Dunn’s multiple comparison correction in (D). ns, not significant; \**P* < 0.05; \*\**P* < 0.01; \*\*\*\**P* < 0.0001, #*P* < 0.05; ###*P* < 0.001. Comparisons in (D) are with WT (*) and syt7KO (#) conditions; full statistics are provided in Data S1.

### STED super-resolution microscopy reveals active zone clustering of all three syt7 isoforms

For syt7 to directly mediate STP by controlling SV dynamics, it must be present in active zones, where release occurs. So, we examined the localization of all three splice variants of syt7 and endogenous syt1 using three-dimensional stimulated emission depletion (STED) microscopy, along with confocal imaging of endogenous bassoon (a marker for active zones at synaptic boutons). Syt7 isoforms were tagged to circumvent the lack of highly specific antibodies. Representative STED (syt7 in magenta and syt1 in cyan) images, along with bassoon (in orange), indicate that the active zone was decorated with each of the three syt7 isoforms (α-, β-, or γ-syt7-FLAG: left, middle, and right panels) and syt1 (Fig. 5A). Qualitatively, all three isoforms displayed similar organization at synapses. To quantify this, we performed cluster analysis on en face synapses containing both syt7 and syt1 clusters, as described in the Supplementary Methods. The number of syt7 and syt1 clusters per bouton was similar across splice-variant conditions (Fig. 5B, syt1 = 1.60 ± 0.13, α-syt7 = 1.64 ± 0.11, β-syt7 = 1.42 ± 0.08, and γ-syt7 = 1.45 ± 0.08, as mean ± SEM). The three variants also showed a similar cluster colocalization with syt1 (Fig. 5C, α-syt7 = 0.98 ± 0.11, β-syt7 = 0.89 ± 0.1, and γ-syt7 = 1.0 ± 0.09, as mean ± SEM). Because the three isoforms exhibit distinct self-association properties, we next asked whether these differences were associated with altered localization to bassoon-positive synapses. All three isoforms were detected at comparable fractions of bassoon-positive puncta (Fig. 5D; α-syt7 = 66%, β-syt7 = 71%, and γ-syt7 = 64%). Thus, despite their distinct biochemical properties, α-, β-, and γ-syt7 show broadly similar active zone-associated localization and clustering, arguing against gross splice-variant-specific differences in synaptic localization.

**Figure 5.**
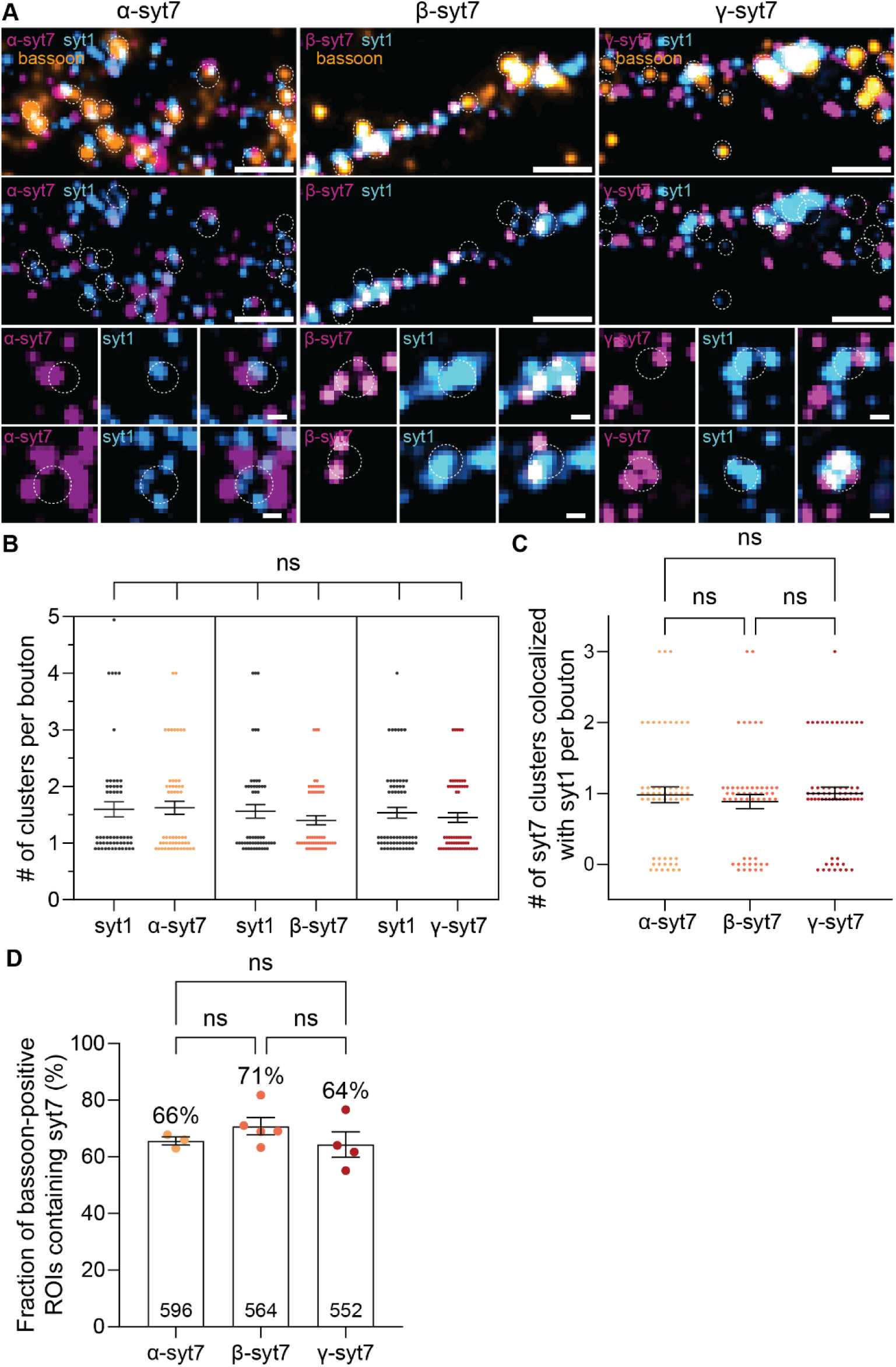
STED super-resolution microscopy shows that clusters of α-, β-, or γ-syt7 colocalize with syt1 clusters in active zones. (**A**) Representative fluorescence images of maximum Z-projections of syt7 (magenta, STED), syt1 (cyan, STED), and bassoon (orange, confocal) in syt7KO mouse hippocampal neurons overexpressing α-, β-, or γ-syt7-FLAG (left, middle, and right, respectively). Dotted white circle indicates bassoon-positive active zones within boutons. Middle panel shows STED images of the three syt7 isoforms, along with syt1. Bottom panel shows two zoomed-in en face views of synapses, visualizing the three syt7 isoforms, syt1, and their overlap. Scale bar, 1 µm (top and middle), 0.1 µm (bottom). (**B**) Frequency of observed clusters of the three syt7 isoforms and syt1 (N = 3, n = 55, 53, and 62 for α-, β-, and γ-syt7 conditions, respectively). The number of clusters was similar across conditions. (**C**) The frequency of colocalized clusters of the three syt7 variants with syt1 at synapses was similar, with a mean of ∼1. Note for (B,C), the en face view synapses that contained both syt7 and syt1 clusters were analyzed. Cluster counts are discrete integers; data points were slightly offset to better visualize overlapping values. (**D**) %Occurrence of the three isoforms of syt7 with bassoon, using 3 independent cultures; the number of puncta is provided in the bar graph. To determine statistical significance, one-way analysis of variance (ANOVA) with Kruskal-Wallis test with Dunn’s multiple comparison correction was used in (B,C), and Tukey’s multiple comparison test with a single pooled variance was used in (D). ns, not significant. Full statistics are provided in Data S1.

### Two-color MINFLUX super-resolution imaging reveals nanometer-scale organization of syt7 clusters at the active zone

While the three isoforms localize and form clusters at the synapse, we next delved into the precise nanometer-scale localization of the α variant, which is thought to be the most abundant isoform^32^, using 3D minimal photon flux (MINFLUX) super-resolution imaging, capable of resolving individual fluorophores at distances of 2-3 nm^34,35,40^. MINFLUX precisely controls the bottle-beam-shaped excitation and achieves nanometer-level resolution by iteratively narrowing the probing pattern to pinpoint the location of fluorescent molecules (Fig. 6A,B). Here, we harnessed a stochastic switching strategy for fluorophores (Abberior FLUX 640 and 680 secondary antibodies for labeling primary syt1 and FLAG antibodies to detect syt1 and α-syt7fl-FLAG), which transitions between fluorescent and non-fluorescent states using reducing buffer conditions, as detailed in the Methods. Due to the complexity of these experiments, MINFLUX imaging of the β and γ variants will be addressed in future studies, in order to determine whether changes in their linker segments affect their nanoscale localization at the active zone.

**Figure 6.**
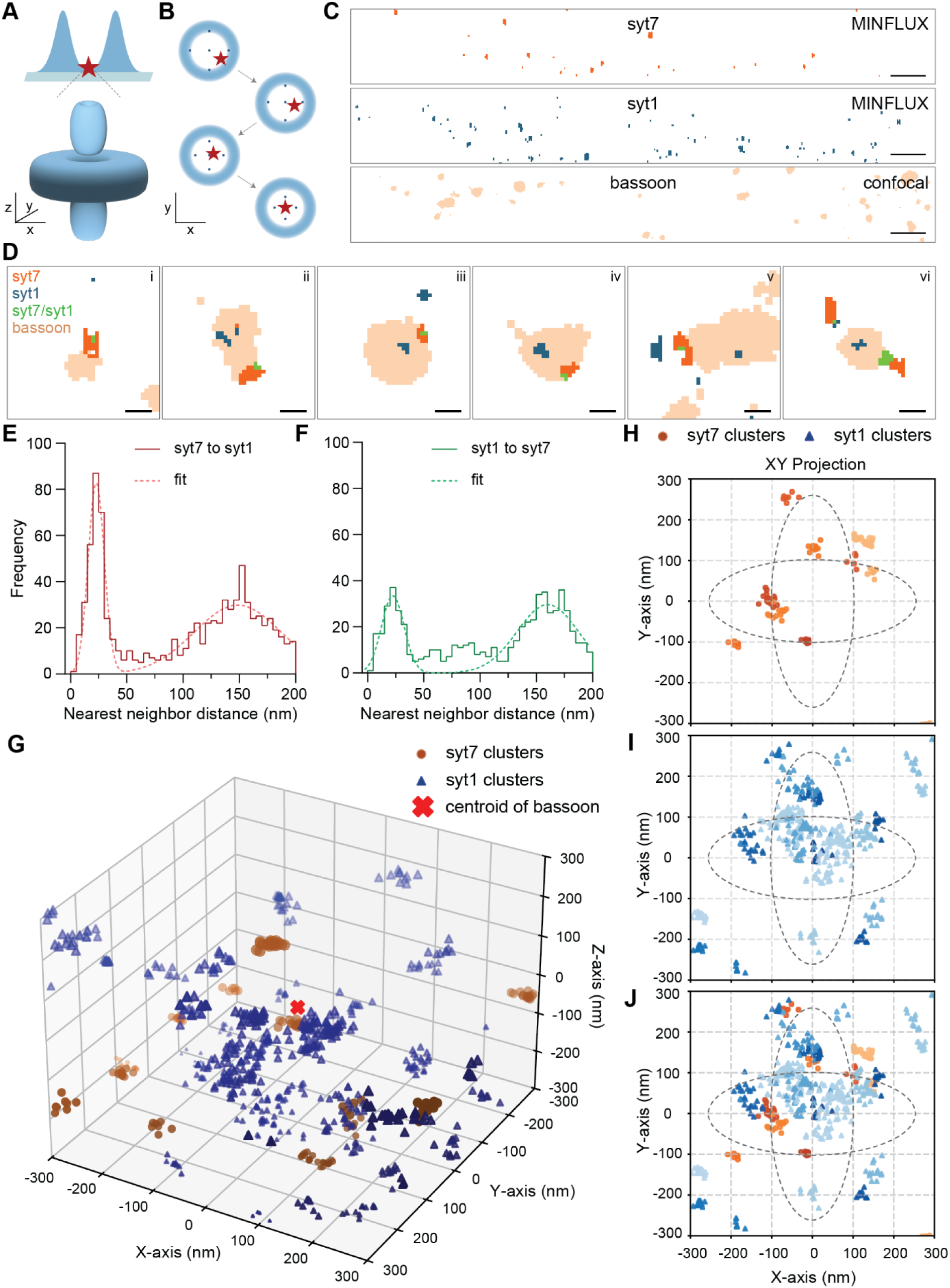
Two-color MINFLUX super-resolution microscopy reveals nanometer-scale organization of syt7 and syt1 at boutons in hippocampal neurons. (**A**) Illustration of a fluorophore (indicated as a red star) excited by a 3D donut beam (or bottle-beam shape; in blue) in minimal photon flux (MINFLUX) super-resolution microscopy. (**B**) 2D schematic of the localization of a fluorophore by a 3D excitation beam progressively narrowing the probing pattern (illustrated by small blue dots) to precisely localize the fluorescent molecule. (**C**) Representative images of maximum Z-projection of syt7 (orange, MINFLUX), syt1 (blue, MINFLUX), and bassoon (tan, confocal) from a FOV. Scale bar, 2 µm. (**D**) Zoomed-in merged images (i – vi; syt7, syt1, and bassoon) from (C); overlap of syt7 and syt1 is represented in green. Scale bar, 0.2 µm. (**E,F**) Histogram of nearest neighbor distances (NND) of syt7 to syt1, and syt1 to syt7, respectively. Both graphs showed a peak around 22 nm, with the syt7 to syt1 peak being ∼2.5 times larger than the syt1 to syt7 NND peak. The broad peak, centered around ∼150 nm in both plots, occurs for any two random fluorescent molecules imaged by MINFLUX. The data were well-fitted with two Gaussian functions (dotted lines). (**G**) 3D representation of syt7 (orange circles) and syt1 (blue triangles) clusters distributed around the bassoon centroid (red cross). Darker colors indicate positive values along the X and Y axes, and larger symbols indicate positive values along the Z axis. (**H-J**) 2D-scatter plot generated from 3D graph in (G), showing syt7 clusters, syt1 clusters, and their overlap projected along the XY axes, respectively. Projections along XZ, and YZ axes are shown in Supplementary Fig. 17A-C. Protein clusters were grouped using different shades. Center (0,0) indicates the bassoon centroid obtained from confocal imaging. Dotted ellipses indicate average size of an active zone, with major and minor axes of ∼500 nm and ∼200 nm, respectively. N=15 FOVs, across six independent neuronal cultures.

We successfully resolved 2-color 3D MINFLUX of syt7 (orange) and syt1 (blue), and achieved a fluorophore localization precision of 5.39 ± 0.26, 5.14 ± 0.26, and 3.01 ± 0.22 nm along the X, Y, and Z axes, respectively (Fig. 6C and Supplementary Fig. 15A,C,D). However, the IgGs used to detect syt7 and syt1 have a linkage error of ∼20 nm ^41^, so the practical localization, which is the quadratic sum of this linkage error plus the instrument precision, was ∼21 nm in the X-Y coordinate. In parallel, we also performed confocal imaging of bassoon (tan) (Fig. 6C). Representative colocalization images of syt7 and syt1 (MINFLUX) with bassoon (confocal) are shown in Fig. 6D. As a negative control, non-transduced syt7KO neurons, stained with the anti-FLAG antibody, exhibited a minimal background signal (Supplementary Fig. 16A). We also note that the syt1 IgG (DSHB, mAB48) has been extensively characterized and does not yield significant background signals using syt1 KO neurons^17,30,42^.

After filtering the 3D data based on localization intensity (Supplementary Fig. 15B,C), and combining data across multiple experiments, we generated frequency distribution plots of the nearest neighbor distances (NND) of syt7 to syt1 and syt1 to syt7, where two peaks are apparent (red and green, respectively, Fig. 6E,F, Supplementary Fig. 15D,E); the dotted curve represents two peaks fitted with Gaussian functions. The small NND peak was at ∼22 nm, and the larger one was centered at ∼150-160 nm (see Supplementary Table 5). The small peak was significantly larger when measuring the distance from syt7 to the nearest syt1 molecule. This suggests that syt1 and syt7 form separate groups, but a few syt1 molecules are consistently clustered very close to the syt7 group. We note that 20% of syt1 is localized to the neuronal surface, as shown by STED microscopy, and pHlourin-syt1 imaging^38,43^, yielding a ∼20% error in our NND calculations.

Next, we sought to localize syt7 and syt1, with respect to a bassoon centroid, as determined via confocal images of the bassoon. Briefly, we calculated the relative vector distances of syt7 and syt1 in X, Y, and Z coordinates from the bassoon centroid, and combined multiple bassoon centroids from multiple images. The data were further processed to find syt7 and syt1 clusters using the DBSCAN algorithm, and presented as a 3D graph (Fig. 6G, see Methods). This plot shows orange circles and blue triangles to indicate syt7 and syt1 clusters, respectively, while a red cross at (0,0,0) denotes the bassoon centroid. The graph is coded such that positive X-Y coordinates are indicated by darker color shades, while positive Z coordinates are denoted by larger symbols. To gain insights into the distribution of syt7 and syt1 around the bassoon center, we plotted 2D projections in XY, XZ, and YZ axes of syt7 (orange circles), syt1 (blue triangles), and both syt7 and syt1 (XY: Fig. 6H-J; XZ, YZ: Supplementary Fig. 17A-C). Dotted black lines indicate the average size of the active zone (major and minor axes diameters of ∼500 nm and ∼200 nm; Fig. 6H-J), isotropically arranged in the XY projection, estimated from electron micrographs and super-resolution imaging^44,45^. Interestingly, syt7 clusters were mainly found at the boundary of the active zone, while no such pattern was observed for syt1. The distribution pattern of syt7 indicates that eleven out of twelve clusters were found within the active zone boundary in the 2D projection, while one was found at the periphery (Fig. 6H); complementing previously observed localization of the syt7 using STORM super-resolution microscopy^46^. In the case of syt1 clusters, which are present on SVs (∼80%) and the axonal plasma membrane (∼20%), we found overlap with the center of the bassoon (at (0, 0) coordinate), and at the edges of the active zone (Fig. 6I). This suggests that syt1 is mainly found at the center of boutons in SV pools, while a ∼20-30% proportion of these clusters were found near the active zone. A merged map of the syt7 and syt1 clusters is shown in Fig. 6J, further illustrating their partial colocalization. Collectively, these findings establish that α-syt7 is mainly found within the boundary of the active zone and α-syt7 and syt1 form distinct clusters, with a proportion of syt1 localizing with syt7 clusters.

## Discussion

Syt7 is a widely expressed^47^, high-affinity calcium sensor crucial for STP^48^. Genebank data and early studies suggested that *syt7* undergoes alternative splicing at the jxm linker^32,36,49^, potentially existing as twelve different isoforms in humans (NCBI Gene ID: 9066). Fukuda et al. established the expression of three splice variants of syt7 in mice: α, β, and γ, which are well conserved in humans, so these are the isoforms investigated in the current study^32^. Here, we demonstrate that alternative splicing of syt7 regulates its self-association and show that these properties correlate with differences in syt7-mediated STP.

Firstly, we discovered that the α and β isoforms of syt7 both underwent liquid-liquid phase separation (LLPS) (Fig. 1D-F), with the β variant showing a slightly higher propensity to form droplets than α (Supplementary Fig. 1A,B). In stark contrast, the γ isoform formed insoluble aggregates (Fig. 1D,G). This difference was conserved in cellular environments; namely, on the plasma membrane of HEK293T cells (Fig. 2A-D, Supplementary Fig. 6A,B) and in the interbouton regions of neurons (Fig. 2E,G-I, Supplementary Fig. 7A,B). The α and β isoforms formed clusters that recovered rapidly after photobleaching (Fig. 2A-C,E,G,H, Supplementary Fig. 7A), whereas the γ isoform showed limited recovery (Fig. 2D,I, Supplementary Fig. 7B). The α- and β-syt7 patches were sensitive to 1,6-HD, consistent with assembly through LLPS (Fig. 2A-I, Supplementary Figs. 6A, 7A). Strikingly, none of the isoforms recovered from photobleaching within presynaptic boutons (Fig. 2F,J-L, Supplementary Fig. 7A,B), suggesting that syt7 is stably anchored, likely through interactions with the presynaptic cytoskeleton^50^. Together, the HEK293T data show that the three full-length syt7 splice variants exhibit distinct mobilities in the plasma membrane of heterologous cells.

Importantly, the same qualitative splice-dependent behavior was observed in hippocampal neurons within inter-bouton regions. These findings suggest that the differences in mobility between the variants are intrinsic. The exception occurs within boutons, where all three variants are immobile, presumably because they bind cytoskeletal elements. Thus, the diffusion of syt7 depends on both alternative splicing and subcellular localization in hippocampal neurons.

The founding member of the syt family, syt1, was also recently shown to undergo LLPS^29^. Both isoforms, syt1 and syt7, sense Ca^2+^ ^23^, but the former is a fast, low-affinity sensor^24^ that triggers SV exocytosis^16–20^, while the latter is a slow, high-affinity sensor^24^ that regulates transient SV docking^25^. In the case of syt1, LLPS separation was shown to play a crucial role in clamping spontaneous SV release, as well as triggering rapid, efficient evoked exocytosis^30^. Droplets formed by syt1, α-syt7, and β-syt7 recover from FRAP to similar extents (Fig. 1H), and the cytoplasmic domains of all three proteins formed droplets in neurons (Supplementary Fig. 4A,B)^29^. However, Ca^2+^ promoted syt1 droplet formation^29^ but had no effect on syt7 droplets (Fig. 1K). The basis for this difference is unclear, but it suggests a form of regulation of syt1 that does not occur with syt7. Rather, syt7 LLPS is regulated by alternative splicing.

The isoform-specific biophysical differences regarding LLPS exhibited by the syt7 splice variants were correlated with distinct functional outcomes for synaptic transmission. Presynaptic function was monitored using iGluSnFR and revealed that the β isoform exhibited enhanced PPF relative to the α isoform (Fig. 3A,B), without any changes in the first response (Supplementary Fig. 10A). Moreover, β rescued the SVRR (Fig. 4C,D) while counteracting synaptic depression more effectively than any of the other rescue conditions, and even better than WT neurons (Fig. 4A,B). Surprisingly, γ, when expressed at the same level as α and β, failed to rescue the SVRR or synaptic depression phenotypes (Fig. 4A-D), yet gave rise to the highest PPF ratio (Fig. 3A,B), with a caveat: it suppressed the initial response (Supplementary Fig. 10A), by decreasing the number of synapses that responded to the first AP (Fig. 3C,D). This shift in release success probability (P_success_), which is a function of n (number of fusion-competent vesicles) and p (fusion probability)^51^, does not appear to significantly change in any other condition except α, which slightly reduced the magnitude of the first response as compared to WT (Fig. 3A,B, Supplementary Fig. 10A). The α variant also enhanced the amplitude of the second response to rescue PPF to WT levels (Fig. 3A,B, Supplementary Fig. 10B), but, like γ, α failed to rescue the SVRR or synaptic depression phenotypes in the KO (Fig. 4A-D). In the context of the α and γ variants findings, we note that syt7KO neurons trend toward increased P_success_ (Fig. 3A,B, Supplementary Fig. 10A)^21,25^. While this trend has not reached significance in the current (Supplementary Fig. 10A) or previous studies^21,25^, it suggests that splice variants that reduce the first response are likely to be expressed at low levels in hippocampal neurons. While this study focused on quantifying presynaptic glutamate release, in future studies, it will be important to assess synaptic transmission, as not all released glutamate necessarily activates postsynaptic receptors and may instead engage presynaptic targets or potentially even glial cells. Thus, comparing iGluSnFR-based release measurements with voltage-clamp recordings could potentially provide further insights regarding the relationship between secretion and functional transmission. In addition, syt7 has been implicated in asynchronous release, but this is difficult to measure accurately with iGluSnFR. Future electrophysiological experiments will provide a quantitative measure of whether alternative splicing affects this slow component of transmission.

For syt7 to function presynaptically to control STP, we sought to determine whether it is present at the active zone, where SVs dock and fuse. Using STED super-resolution microscopy, we found that all three syt7 splice variants formed clusters within bassoon-positive active zone and showed similar colocalization with endogenous syt1 (Fig. 5A-D). This indicates that the functional differences among the α-, β-, and γ-syt7 variants are unlikely to arise from gross differences in synaptic localization. Using MINFLUX super-resolution imaging, we further localized α-syt7 to clusters within the active zone boundary (Fig. 6H). Some of these clusters overlapped with syt1 clusters (Fig. 6I,J), consistent with docked syt1-bearing SVs colocalizing with syt7 at the active zone (Supplementary Fig. 18A). Hence, syt7 is well-positioned to regulate STP by controlling aspects of the SV, and in particular, the docking step, as discussed below^21,25^.

We emphasize that syt7 is not tailored to trigger SV release but rather modulates synaptic strength in response to recent activity. Owing to its higher Ca^2+^ affinity and slower kinetics, syt7 is particularly sensitive to residual Ca^2+^ ^3,52^, which lingers in the presynaptic terminal after an AP. Recently, syt7 has been proposed to regulate transmission by mediating activity-dependent “transient” SV docking^21,25^. We further explored this idea and interrogated whether SVs bind directly to syt7. Indeed, HaloTag-pull down assays indicated an interaction between α-syt7cyto and purified SVs, and this interaction was weakly promoted by Ca^2+^ (Ca^2+^/EGTA condition: 1.34 ± 0.23; Supplementary Fig. 19A,B). We further addressed this docking hypothesis by exploring binding partners of syt7 via immunoprecipitation (IP; Supplementary Fig. 19C,D) using rat hippocampal neurons expressing α-syt7-fl-FLAG. In the absence of cross-linkers, we identified a sole and specific interaction between syt7 and syt1 (Supplementary Fig. 19D), as observed previously^28^. We validated these findings using immobilized syt7 and syt1-fl that had been reconstituted into liposomes and, again, observed an interaction that is modestly enhanced by Ca^2+^ (Ca^2+^/EGTA: 1.46 ± 0.15; Supplementary Fig. 19A,B). It is likely that the weak Ca^2+^-dependence is mediated by the C2-domains of one or both of these proteins, but, interestingly, we also found that the jxm linkers of syt7 and syt1 directly bind to one another in a concentration-dependent manner (Supplementary Fig. 19E,F). Moreover, these jxm linker segments mediate LLPS, and both α- and β-syt7cyto coalesce with syt1cyto droplets *in vitro* (Supplementary Fig. 20A,B), and α-syt7cyto coalesces with syt1cyto in neurons (Supplementary Fig. 20C), further confirming their interaction.

Together, the findings reported here are consistent with a model in which syt7 is a docking protein that mediates PPF^21,25^. After a single AP, ∼40% of all SVs undock^53^, and syt7 is crucial for rapidly re-docking vesicles to refill the “slots” for the next round of release^25^, potentially mediated, in part, by coalescence of syt7-syt1 phase-separated surfaces. This would occur during activity, as – again – docking is unaffected under resting conditions in syt7KO neurons^25^, and other known protein and lipid interactions have been shown to participate in the attachment of SVs to the active zone membrane^54,55^. This 2D LLPS interaction, mediated by the syt7 and syt1 jxm linkers (Supplementary Figs. 18E,F, 19A,B), might juxtapose their C2-domains to enhance Ca^2+^-promoted binding between them, thus recruiting more SVs to release sites during ongoing activity. Surprisingly, iGluSnFR measurements revealed that only the β splice variant rescued the SVRR and synaptic depression phenotypes (Fig. 4A-D), while α and γ failed, despite the finding that all three rescued PPF (Fig. 3A,B). Thus, alternative splicing can uncouple the role of syt7 in these two different forms of short-term plasticity. While the leading model for syt7 function in PPF is via activity-dependent docking, the findings reported here reveal that the role of syt7 during synaptic depression is more complex, involving either additional or completely different mechanisms. Hence, differences in the functions of α, β, and γ provide a molecular handle for further investigation of the mechanisms underlying STP. Future zap-and-freeze electron microscopy studies of neurons expressing each splice variant will shed light on the transient docking question. Another important goal is to purify the cytoplasmic domains of all three variants in non-droplet and non-aggregated forms to enable biochemical studies to investigate how the jxm linkers affect Ca^2+^ and effector binding.

In summary, the current study demonstrates that alternative splicing of the syt7 jxm linker serves as a molecular switch, toggling the protein between dynamic, functional condensates and static, potentially disruptive aggregates. These properties correlated, to some extent, with a functional impact on synaptic plasticity and might even play a role in determining the P_success_. Since alternative splicing affects only the linker domain of syt7, it seems likely that differences in linker-mediated self-association underlie the observed differences in function. It will be important to delve deeper into this issue via mutational analysis and linker-swapping experiments, to further establish causation. A particularly challenging issue is to understand how subtle differences in the propensity to phase separate, between the α and β variants, translate to changes in PPF, and to determine how the γ variant, which forms aggregates, is able to sharply suppress the fraction of active synapses responding to a stimulus. In this context, it is notable that the *Drosophila* ortholog of syt7 was reported to be inhibitory^56^, but whether this ortholog phase separates or aggregates has yet to be determined. Given that protein condensation and aggregation are inherently concentration-dependent phenomena, a crucial future direction will be to investigate how lower expression levels of the γ-variant affect its biophysical and functional properties. Furthermore, protein aggregation is a recognized hallmark of several neurodegenerative diseases^57^, so dysregulation of the γ variant can potentially contribute to pathophysiology. Finally, while this study focused on the α, β, and γ isoforms of syt7, it is unclear which mixture of variants is expressed at which synapses. Although syt7 splice variants are developmentally and regionally regulated, the mechanisms controlling alternative splicing, the neuronal subtypes in which each variant is expressed, the relative abundance of each isoform, and whether neuronal activity impacts their expression, remain unknown. Addressing these questions will require spatially resolved transcriptomics, along with cell-type-specific translational profiling. Moreover, we recognize that there may be other, undetected syt7 isoforms expressed in the brain (NCBI Gene ID: 9066). This new direction is crucial, as the three splice variants characterized thus far, as shown here, have distinct biophysical and functional properties. Finally, we note that among the seventeen isoforms of syt (not including splice variants), ten isoforms possess IDRs within their jxm linkers. It will be interesting to determine which isoforms undergo LLPS, and to address how phase separation impacts their function.

## Materials and Methods

### Two-color MINFLUX imaging

Minimal photon flux (MINFLUX) super-resolution imaging setup was done as previously described^34,40^. Briefly, an excitation laser beam (green) is shaped by a vortex-phase mask, forming a donut-shaped intensity spot in the focal plane of the objective lens. Photons emitted by the fluorescent molecule (star) are collected by the objective lens and directed toward a fluorescence bandpass filter and a confocal pinhole by using a dichroic mirror. Intensity modulation and deflection, as well as photon counting, are controlled by a field-programmable gate array. Additional methods are described in the Supplementary Information.

## Acknowledgments

We thank Jessica Matthias, Otto Wirth (Abberior GmbH), and Dr. Nicolas Piguel (Northwestern University) for assisting in super-resolution imaging, Dr. Abbas Rizvi and Yuchen Xu for transcriptomics analysis, the Institute of Genomic Biology, UIUC, for the MINFLUX instrument, the Center for Advanced Microscopy (RRID: SCR_020996), and Northwestern University for the Abberior MIRAVA Polyscope, supported by NCI-CCSG-P30-CA060553 awarded to the Robert H Lurie Comprehensive Cancer Center. Mice used in this study were obtained from the Mary Lyon Center at MRC Harwell (MLC), and the following award is acknowledged: MC_UP_2201/2. The MLC is also a member of the International Mouse Phenotyping Consortium and has received funding from the National Institute for Health (5UM1HG006348-08) for generating and/or phenotyping the C57BL/6N-Syt7^em1(IMPC)H^/H mice. ERC is an Investigator of the Howard Hughes Medical Institute (HHMI). This article is subject to HHMI’s Open Access to Publications policy. HHMI lab heads have previously granted a nonexclusive CC BY 4.0 license to the public and a sublicensable license to HHMI in their research articles. Pursuant to those licenses, the author-accepted manuscript of this article can be made freely available under a CC BY 4.0 license immediately upon publication.

## Funding

NIH grants R01MH061876, R35NS136306 (ERC)

Howard Hughes Medical Institute (ERC)

## Data and materials availability

All materials, blots, and statistical information are available in the supplemental materials.

For raw data and code, Dryad:http://datadryad.org/share/LINK_NOT_FOR_PUBLICATION/BDtrDQSWRIZd XvUczztX6v2f9ldyTiiNUD490ZRUhsE

GitHub: https://github.com/nikunjrmehta/MINFLUX-Data-Analysis.git

## Supplementary Information

Supplementary Figures. 1 to 20

Supplementary Tables 1 to 5

Supporting Material and Methods

Data S1 to S2

## Ethics Statement

Animal care and use in this study were conducted under guidelines set by the NIH’s *Guide for the care and use of laboratory animals* handbook. Protocols were reviewed and approved by the Animal Care and Use Committee at the UW-Madison (Laboratory Animal Welfare Public Health Service Assurance Number: A3368-01).

## Author Contributions

Conceptualization: NM, ERC

Formal analysis: NM, DTL, SM, SK, AJ

Investigation: NM, DTL, MW, SS, SM, AJ,

ERC Methodology: NM, DTL, ERC

Software: NM, DTL, SK, AJ

Visualization: NM, DTL, AJ, ERC

Writing: NM, ERC

