## Supplementary Material for "Alternative splicing of synaptotagmin 7 regulates oligomerization and short-term synaptic plasticity"

**The PDF file includes:**

Supplementary Figs. 1 to 20
Supplementary Tables 1 to 5
Supporting Material and Methods

**Other Supplementary Materials for this manuscript include the following:**

Data S1 and S2

### Supplementary Figures

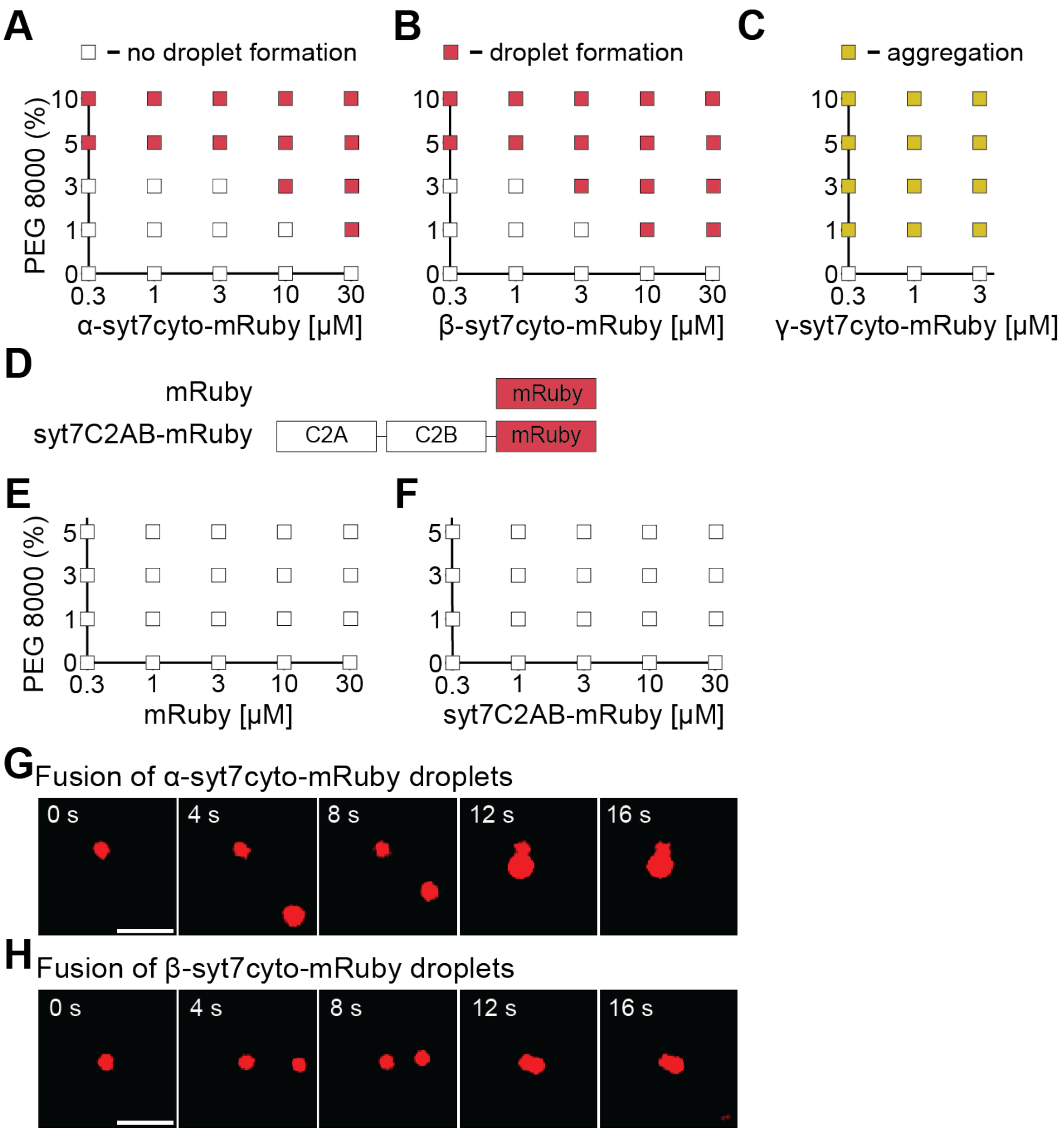

**Supplementary Figure 1. Phase diagram of α, β, and γ-syt7cyto variants and fusion of α and β-syt7cyto droplets.**

(**A-C**) Phase diagram of alternative splice variants of syt7 (α-, β-, and γ-syt7cyto, respectively) at indicated [protein] and [PEG 8000]. Red and yellow squares indicate the formation of droplets and aggregates, respectively, whereas empty squares indicate no effect. Note, the β variant had a slightly higher propensity to form droplets than α-syt7cyto, whereas γ-syt7cyto formed aggregates under all PEG 8000 conditions. (**D**) Control constructs: fluorescent marker, mRuby and syt7C2AB-mRuby. (**E-F**) Phase diagram of the controls in (D) at indicated [protein] and [PEG 8000], with a similar color-scheme as (A-C). No droplets or aggregates were observed for the controls. (**G,H**) Fusion of α- and β-syt7cyto droplets over time. Scale bar, 4 μm (G,H). For all experiments, N≥3 FOVs from three independent trials. The buffer used in (A-C,E,F) was 25 mM Tris-HCl (pH 7.4), 100 mM NaCl, and the indicated % PEG 8000.

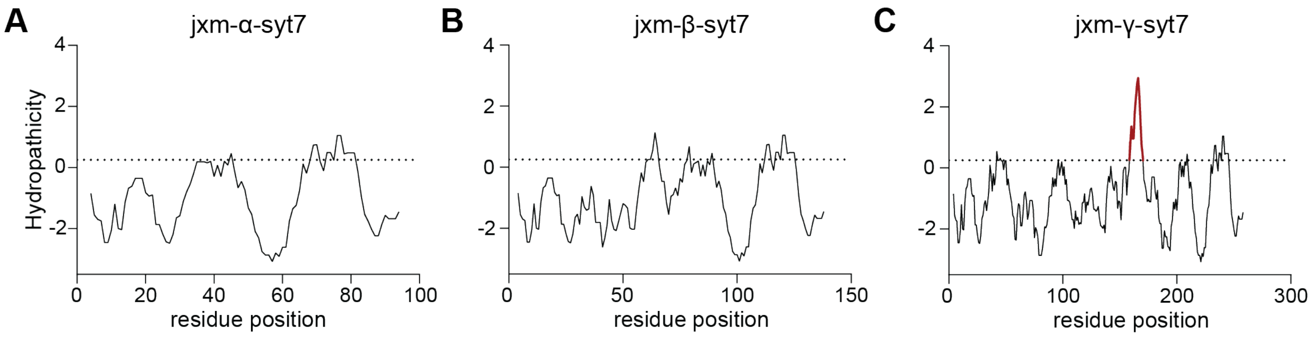

**Supplementary Figure 2. Hydropathicity index of the juxtamembrane linkers of syt7 alternative splice variants.**

(**A-C**) Residue-based hydropathicity score of the juxtamembrane linkers of α-, β-, and γ-syt7. Scores were calculated based on Kyte and Doolittle scoring index. Note, only γ-syt7 had a significant hydropathicity index, indicated in red, for a twelve-residue segment (159- EGRMVVLSLVLG -170). A dashed line at a score of 0.25 was used as a cut-off.

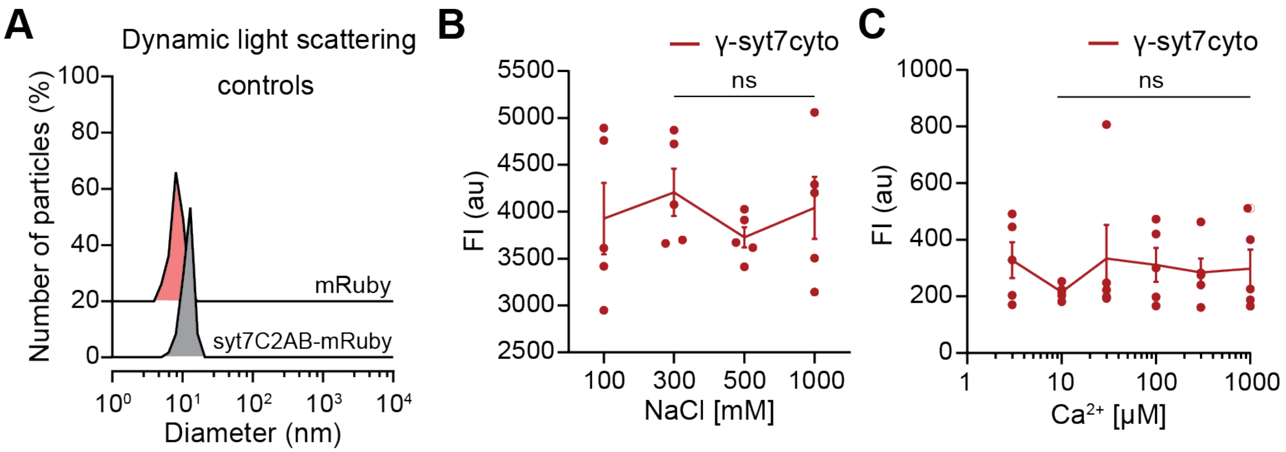

**Supplementary Figure 3. DLS controls and effect of salt and $Ca^{2+}$ on γ-syt7cyto.**
(**A**) Dynamic light scattering (DLS) of the proteins illustrated in Supplementary Fig. 1D, showing number of particles (%) as a function of diameter. Both mRuby and syt7C2AB-mRuby had single peaks corresponding to monomers. (**B,C**) Effect of salt and $Ca^{2+}$ on γ-syt7cyto aggregates, respectively; neither had an effect on the aggregates. Note the buffer used in (B) was 25 mM Tris-HCl (pH 7.4), 100 mM NaCl, and 3% PEG 8000; the same buffer, but lacking PEG 8000, was used in panel (C). N=3; five fields of view (FOVs) were analyzed for each condition tested; data are represented as mean ± SEM. ns indicates not significant.

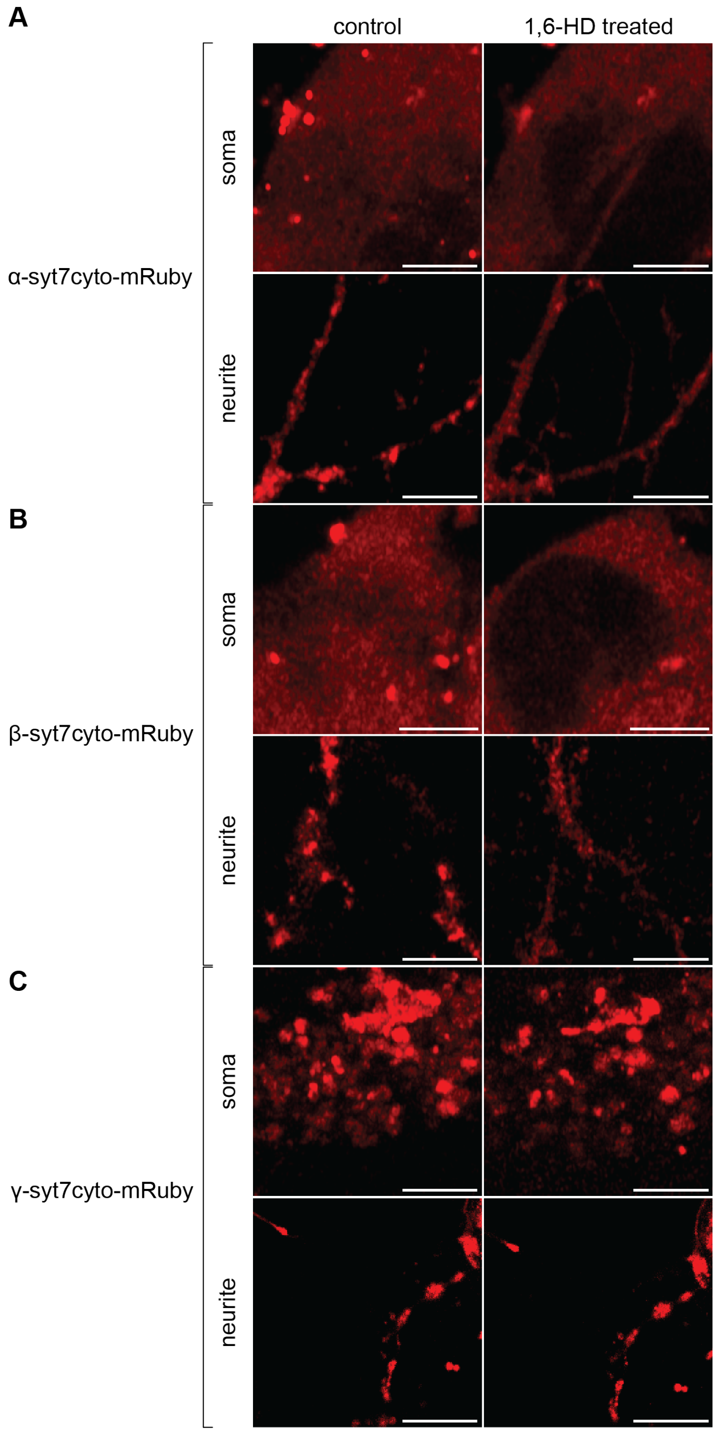

**Supplementary Figure 4. α, β, and γ-syt7cyto form clusters in rat hippocampal neurons.**

(**A-C**) Fluorescence images, captured on a confocal microscope, of the soma and neurites of rat hippocampal neurons transfected with α, β, or γ-syt7cyto-mRuby, respectively, under control and 10% 1,6-HD conditions. Upon 1,6-HD treatment, α- and β-syt7cyto droplets were reduced, while γ-syt7cyto aggregates were unaffected. Scale bar, 4 μm (A-C). N=3 from three independent cultures.

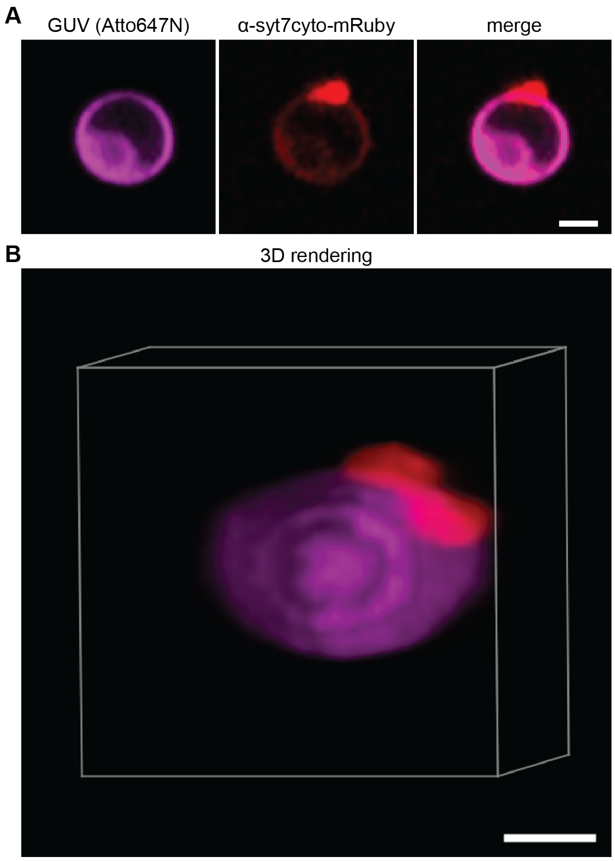

**Supplementary Figure 5. α-syt7 2D liquid-liquid phase separation on the surface of a GUV.**

(**A**) A Z-section plane of an Atto-647-labeled GUV (93.5% DOPC, 5% 18:1 DGS-NTA, 1.5% Atto647N DOPE; magenta) and α-syt7cyto-mRuby (red) captured on a confocal microscope. 2D LLPS of the α-syt7cyto-mRuby 'droplet' on the surface of GUV is apparent. Scale bar, 10 µm. (**B**) 3D rendering of the 2D LLPS of α-syt7cyto-mRuby 'droplet' on the surface of GUV. Scale bar, 2 µm. N=3 independent trials.

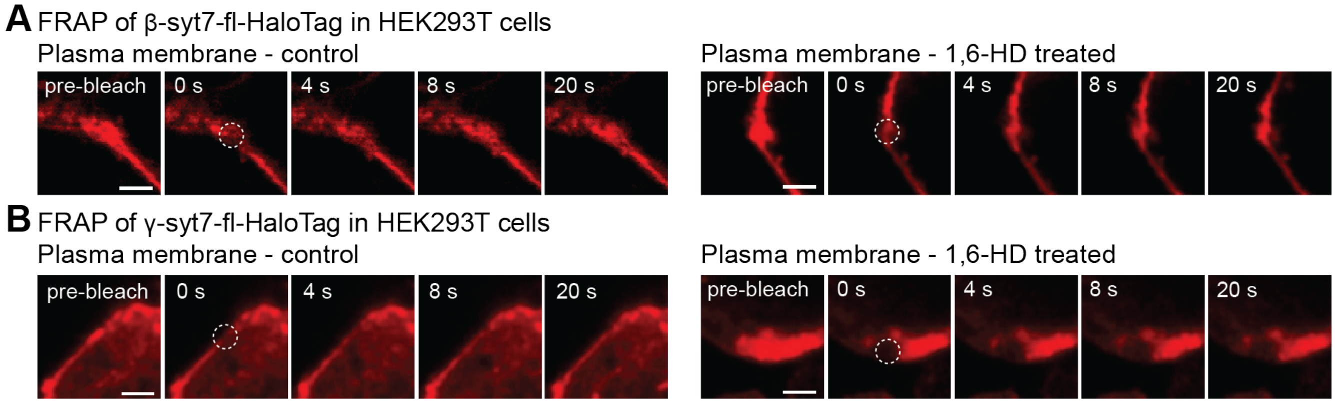

**Supplementary Figure 6. FRAP of β- and γ-syt7-full length on the plasma membrane in HEK293T cells.**

(**A**) Time series of fluorescence recovery after photobleaching (FRAP) at the plasma membrane of HEK293T cells expressing β-syt7-full length(fl)-HaloTag, under control and 10% 1,6-HD conditions, respectively. Scale bar, 2 µm. (**B**) same as (A), but for γ-syt7-fl-HaloTag. Note that β-syt7, but not γ-syt, recovered. Scale bar, 2 µm. N≥10-15 bleached regions across conditions from three independent trials. HaloTag-fusion proteins were labeled with JF549 fluorescent dye.

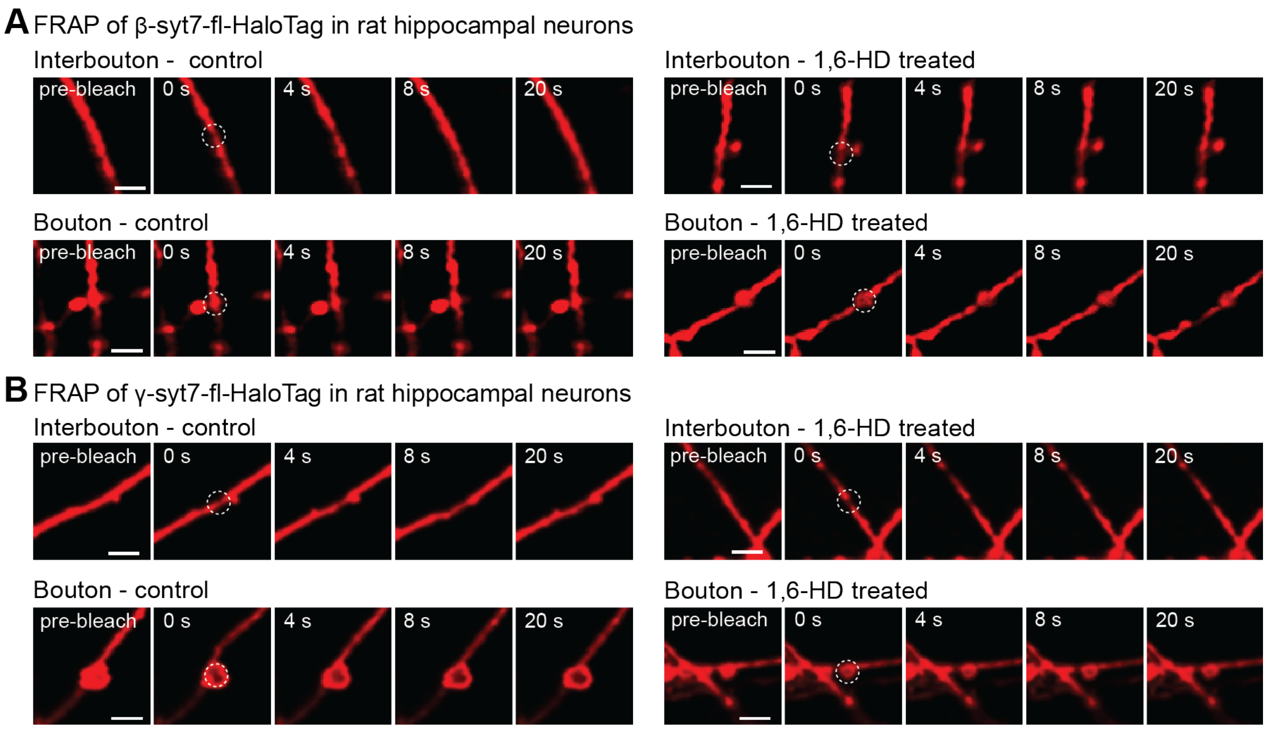

**Supplementary Figure 7. FRAP of β- and γ-syt7-full length in synaptic boutons and interbouton regions of hippocampal neurons.**

(**A**) Time series of FRAP at the indicated locations in rat hippocampal neurons expressing β-syt7-fl-HaloTag, under control and 10% 1,6-HD conditions. Scale bar, 2 μm. (**B**) same as (A), but for γ-syt7-fl-HaloTag. Scale bar, 2 μm. Neither isoform recovered from FRAP in boutons, while β-syt7, but not γ-syt7, recovered within the interbouton regions. Scale bar, 2 μm. N≥10-15 bleached regions across conditions from three independent trials. HaloTag-fusion proteins were labeled with JF549 fluorescent dye.

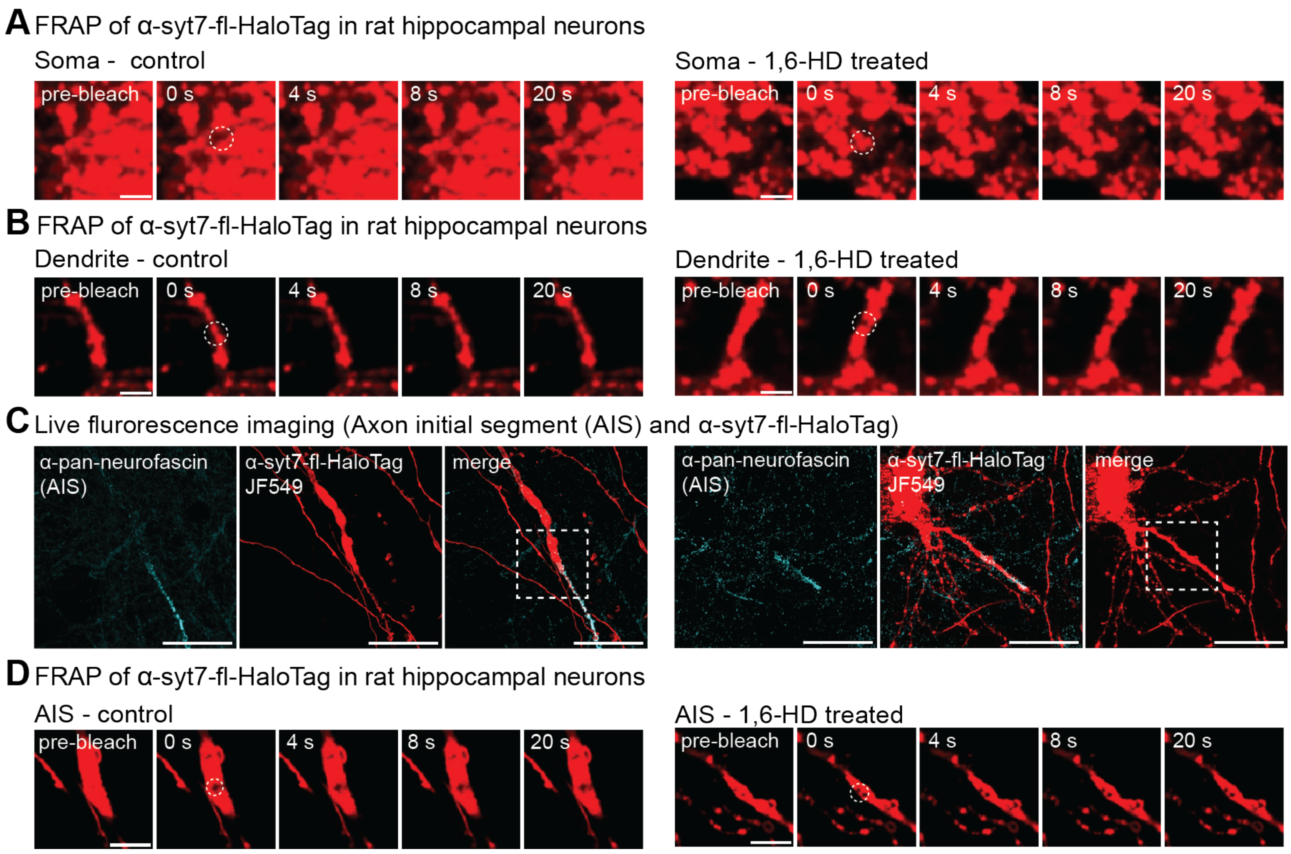

**Supplementary Figure 8. FRAP of α-syt7-full length in the soma, dendrites, and AIS of hippocampal neurons.**

(**A,B,D**) Time series of FRAP at the soma, dendrite, and axon initial segment (AIS), respectively, under control and 1,6-HD conditions. While syt7 is mainly an axonal plasma membrane protein in neurons, overexpression results in spill-over into the somato-dendritic compartment. Surprisingly, this mistargeted protein is relatively immobile, as the bleached area recovered less than 15% across all conditions. Scale bar, 2 μm (A,B), 5 μm (D). (**C**) Live cell imaging of rat hippocampal neurons expressing α-syt7-fl-HaloTag, visualized using JF549 fluorescent dye. AIS was marked extracellularly on live cells with an anti-pan-neurofascin antibody, followed by labeling with an AF647 secondary antibody. Scale bar, 20 μm. N≥10-15 bleached regions across conditions from three independent trials. Box indicated the FOV used for bleaching in (D).

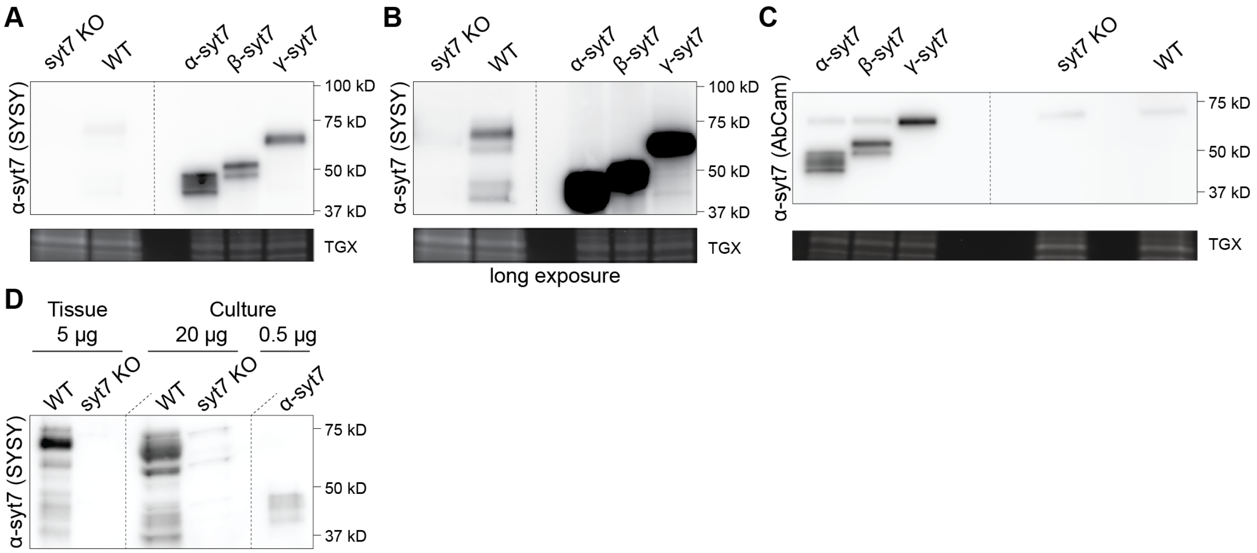

**Supplementary Figure 9. Immunoblotting of syt7 splice variants.**

(**A**) Representative Western blot from 15 days *in vitro* (DIV) lysates from cultured mouse hippocampal neurons showing syt7KO (CRE-treated), WT, and syt7KOs expressing α, β, or γ-syt7, along with a loading control, imaged using stain-free method (indicated as TGX, tris-glycine extended method, BioRad). Probing with an anti-syt7 antibody (SYSY) confirmed KO of syt7, but the syt7 band was weak. (**B**) With long exposure, the same blot clearly shows KO of syt7. SYSY antibody is sensitive to the juxtamembrane linker and thus recognizes the alternative splice variants differentially. (**C**) Probing with anti-syt7 (AbCam) antibody, which specifically recognizes the C2 domains, and does not differentiate between the alternative splice variants, shows expression of the three splice variants, with equal loading in (A). Note that the AbCam antibody is not sensitive enough to detect syt7 WT levels. (**D**) Representative immunoblot of WT and syt7KO from: hippocampal tissue from P15 mice, DIV15 hippocampal cultures; also, syt7KO cultures expressing α-syt7, loaded with 5 (tissue), 20 (culture), and 0.5 μg (culture) of total protein, respectively, and probed with an anti-syt7 antibody (SYSY). Densitometry revealed that α-syt7 (~45 kD) is downregulated in cultured neurons ~3.5-fold as compared to brain tissue, and α-syt7 rescue is expressed ~60 and ~16-fold over cultured neurons and brain tissue, respectively. With equal protein loading, (C) shows β- and γ-syt7 were expressed at ~0.9 and ~0.8 expression levels of α-syt7. N≥3 blots, with three independent cultures.

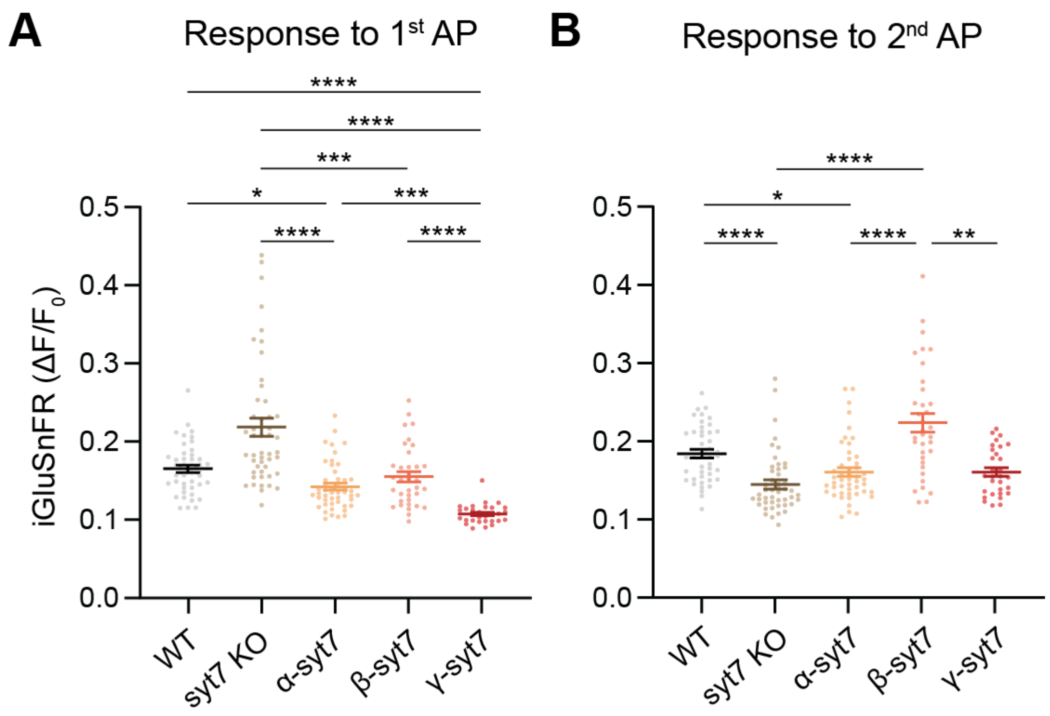

**Supplementary Figure 10. iGluSnFR signal in response to the first and second action potentials.**

(**A,B**) Quantification of peak iGluSnFR first and second responses ($\Delta F/F_0$), respectively, in the first 10 ms bin after applying two APs separated by 50 ms. As compared to the WT condition, the responses to the first stimulus were significantly lower for α- and γ-syt7. Number of FOVs analyzed: 44, 48, 48, 35, and 30 for WT, syt7KO, α-, β-, or γ-syt7 conditions, respectively, across three or more independent culture preparations; data are represented as mean ± SEM. To determine statistical significance, one-way analysis of variance (ANOVA) with Kruskal-Wallis test with Dunn's multiple comparison correction was used in (A,B). *$P < 0.05$; **$P < 0.01$; ***$P < 0.001$; ****$P < 0.0001$. Full statistics are provided in Data S2.

A

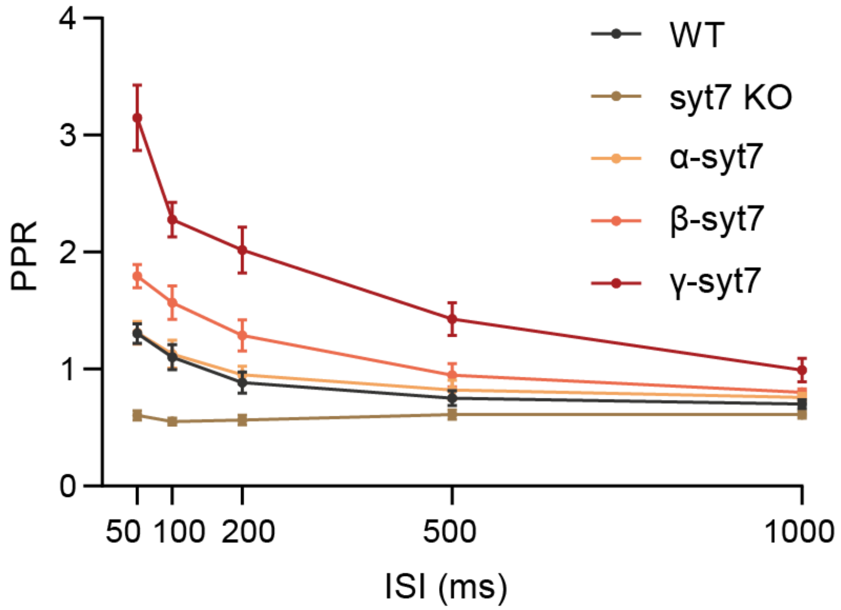

**Supplementary Figure 11. Dependence of paired-pulse facilitation on inter-stimulation intervals of the three syt7 variants.**

(**A**) Paired-pulse ratio (PPR) measured across five inter-stimulation intervals (ISIs) of 50, 100, 200, 500, and 1000 ms. Data are shown as mean ± 95% CI; n ≥ 26 cells from three independent experiments. Statistical annotations are omitted for clarity. To determine statistical significance, a two-way analysis of variance (ANOVA) with Tukey's multiple comparisons test and a single pooled variance was used in (A). Full statistics are provided in Data S2. Curves were fitted with either single- or double-exponential decay, and the results were reported in Supplementary Table 4.

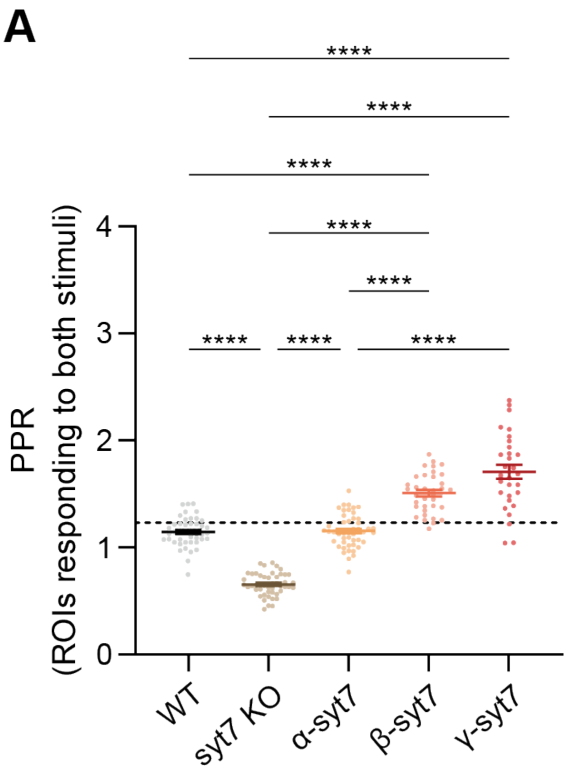

**Supplementary Figure 12. iGluSnFR PPR analysis for boutons that responded to both APs.**

(**A**) Quantification of paired-pulse ratio (PPR) of peak iGluSnFR responses ($\Delta F/F_0$) from ROIs responding to both first and second stimuli at 20 Hz. Dashed line indicates WT PPR from Fig. 3B. Note: a similar trend is observed as Fig. 3B. PPRs calculated from ROIs responding to both stimuli contribute 94, 95, 92, 90, and 71% to the combined PPR of the five conditions tested. Number of FOVs analyzed: 44, 48, 48, 35, and 30 for WT, syt7KO, α-, β-, or γ-syt7 conditions, respectively, across three or more independent culture preparations; data are represented as mean ± SEM. To determine statistical significance, one-way analysis of variance (ANOVA) with the Kruskal-Wallis test and Dunn's multiple-comparison correction was used in (A). *$P$ < 0.05; **$P$ < 0.01; ***$P$ < 0.001; ****$P$ < 0.0001. Full statistics are provided in Data S2.

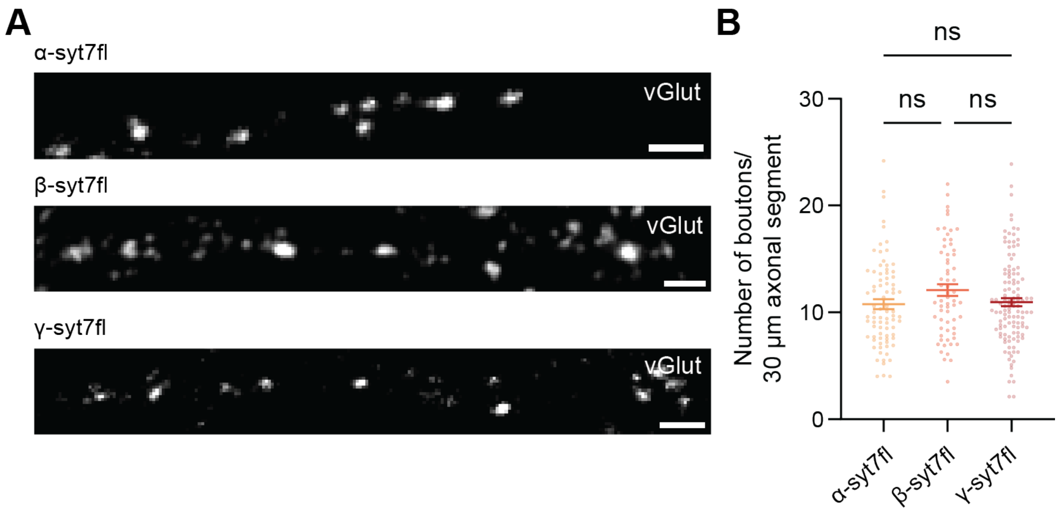

**Supplementary Figure 13. A similar number of vGlut-positive synapses in syt7KO neurons expressing each of the three splice variants of syt7.**

(**A,B**) Representative fluorescence images and quantification of the number of total vGlut-positive synapses in syt7KO hippocampal neurons expressing α, β, or γ-syt7. Neurons were stained with a vGlut antibody, and the number of puncta was quantified from multiple 30 µm axonal segments across conditions. Scale bar, 2 µm. N=10 FOVs for each condition from two culture preparations. To determine statistical significance, one-way analysis of variance (ANOVA) with Kruskal-Wallis test with Dunn's multiple comparison correction was used in (B). ns, not significant; *$P < 0.05$; **$P < 0.01$; ***$P < 0.001$; ****$P < 0.0001$. Full statistics are provided in Data S2.

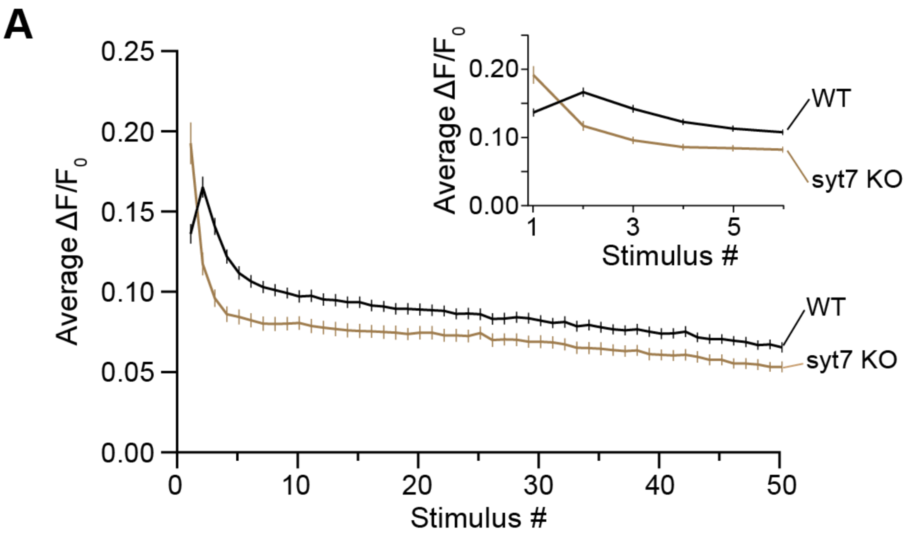

**Supplementary Figure 14. Syt7 counteracts synaptic depression.**

(**A**) For clarity, synaptic depression curves from only WT and syt7KO neurons from Fig. 4B were plotted, to facilitate direct comparisons. The graph shows the average amplitude of iGluSnFR responses ($\Delta F/F_0$) from mouse hippocampal neurons during high frequency stimulation (HFS; 50 action potentials at 20Hz) for WT and syt7KO, shown in black and brown, respectively. Inset shows the first six responses for the two conditions.

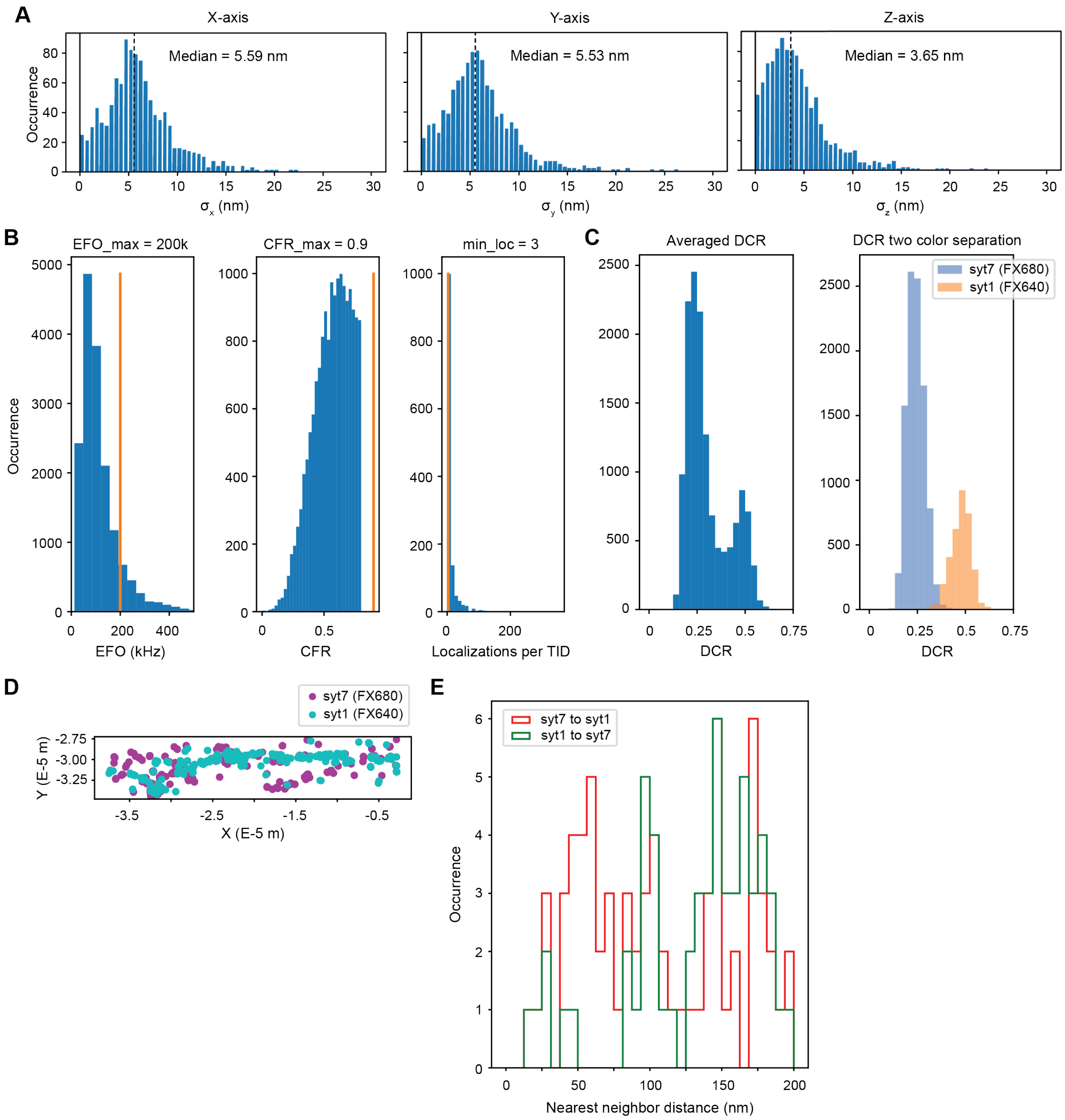

**Supplementary Figure 15. Resolution and filtering of MINFLUX data.**

(**A**) Representative histogram of standard deviations (σ) of localization precision along X, Y, and Z axes. $\sigma_x = 5.39 \pm 0.26$, $\sigma_y = 5.14 \pm 0.26$, and $\sigma_z = 3.01 \pm 0.22$. (**B**) Representative histograms from a 2-color 3D MINFLUX image showing effective frequency at offset (EFO), center frequency ratio (CFR), localization per TIDs (Trace IDs) set as EFO_max = 200k Hz, CFR_max = 0.9, and localization when TID > 3. After filtering the data, (**C**) the detection channel ratio (DCR) was calculated by spectral unmixing of the average DCR plot. The two peaks were fitted with Gaussian functions, indicative of the FX680 (syt7) and FX640 (syt1) dyes, respectively. (**D**) DCR-based color assignment was used to separate FX640 and FX680 in cyan and magenta, and (**E**) nearest neighbor distances were plotted for FX640 to FX680 (indicated as syt7 to syt1; red) and vice-versa (indicated as syt1 to syt7; green), as described in methods. N=15 images from 5 independent culture preparations. Data are represented as mean ± SEM.

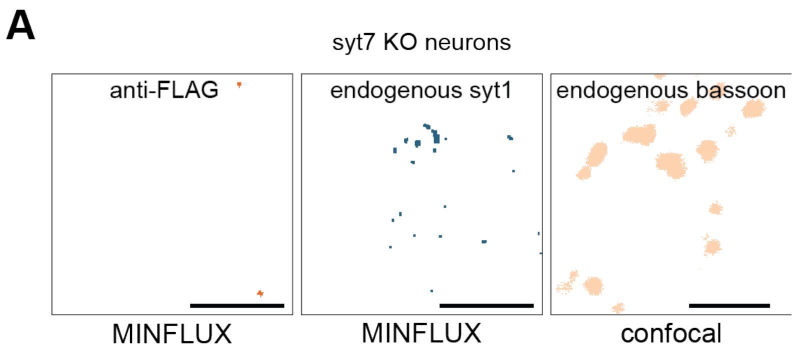

**Supplementary Figure 16. MINFLUX imaging controls: non-transduced syt7KO neurons**

(**A**) Representative images of maximum Z-projection of syt7 (stained with an anti-FLAG antibody in orange, MINFLUX), syt1 (stained with mAb48 in blue, MINFLUX), and bassoon (stained with SYSY 141005 in tan, confocal) from a FOV of non-transduced syt7KO mouse hippocampal neurons. In the absence of transduced FLAG-tagged syt7, little signal was observed using the anti-FLAG antibody. Endogenous syt1 and bassoon expression were normal in the syt7KO neurons. Scale bar, 2 μm.

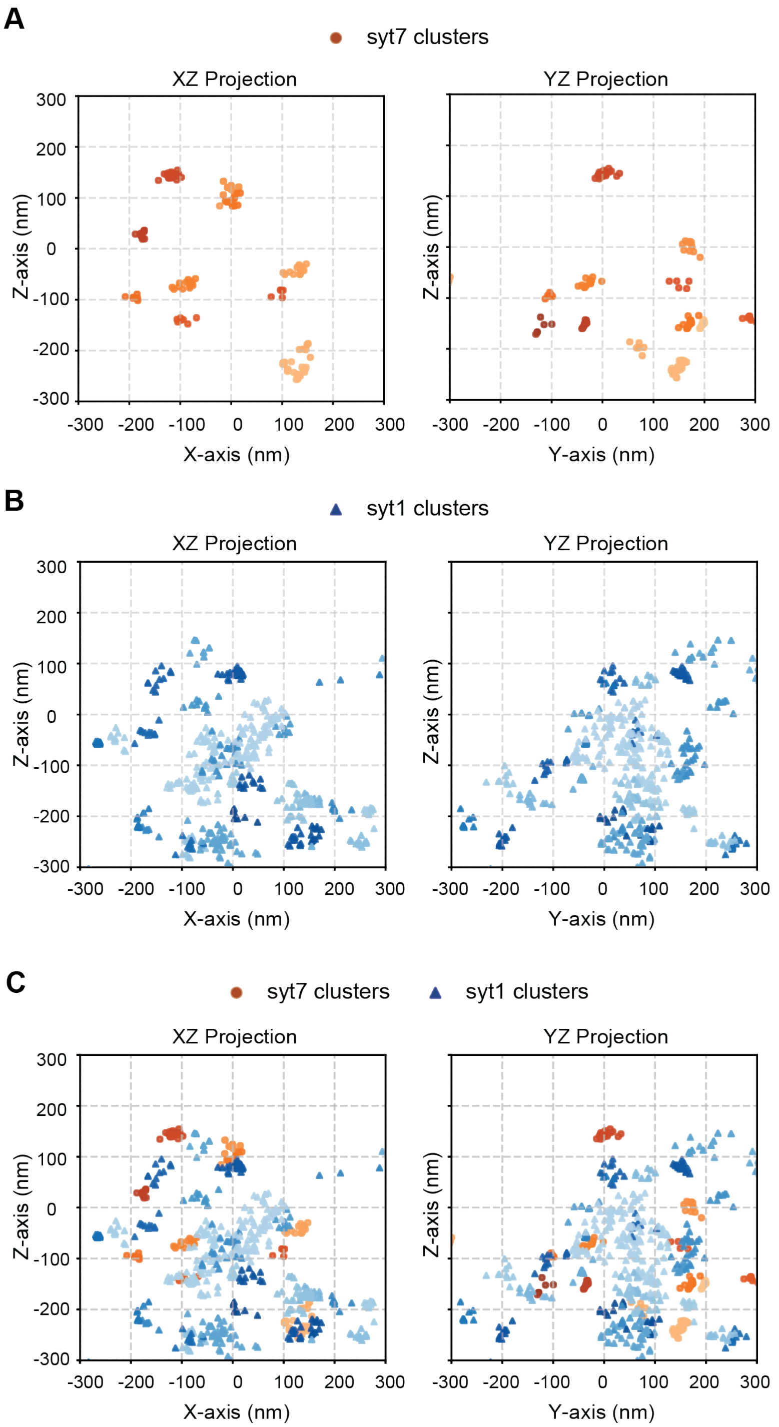

**Supplementary Figure 17. 2D projections from 3D scatter plots of syt7 and syt1 clusters along the XZ and YZ axes.**

(**A-C**) 2D scatter plot generated from the 3D graph shown in Fig. 6G, of syt7 and syt1 clusters, illustrating the projections along the XZ and YZ axes. The center (0,0) indicates the centroid of the bassoon obtained from confocal imaging. Syt7 and syt1 clusters are represented in orange circles and blue triangles, respectively. Note, the clusters are grouped using the same shade of color.

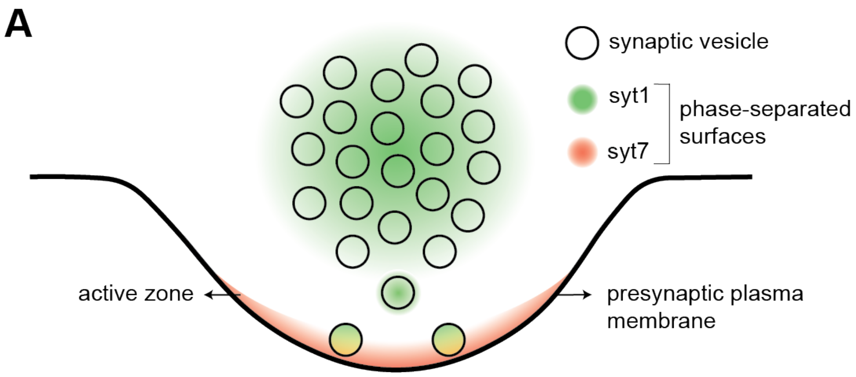

**Supplementary Figure 18. Model describing how syt7-syt1 interactions might contribute to SV docking**

(A) Schematic of SV docking at the active zone of the presynaptic plasma membrane at a bouton. SVs are denoted by circles; syt1 phase-separated surface in green, and syt7 phase-separated surface in red. Syt7-syt1 LLPS 'surfaces' adhere to each other, contributing to another protein-protein interaction in the docking pathway. Since syt7KO neurons do not show defects in docking at steady state[1,2], we propose that the calcium-dependent increase in the avidity of this interaction, likely mediated by the C2-domains[3], contributes to docking reactions during activity. In this model, the interaction of the syt7 and syt1 jxm regions might serve to poise the C2-domains for rapid, efficient interactions during ongoing activity.

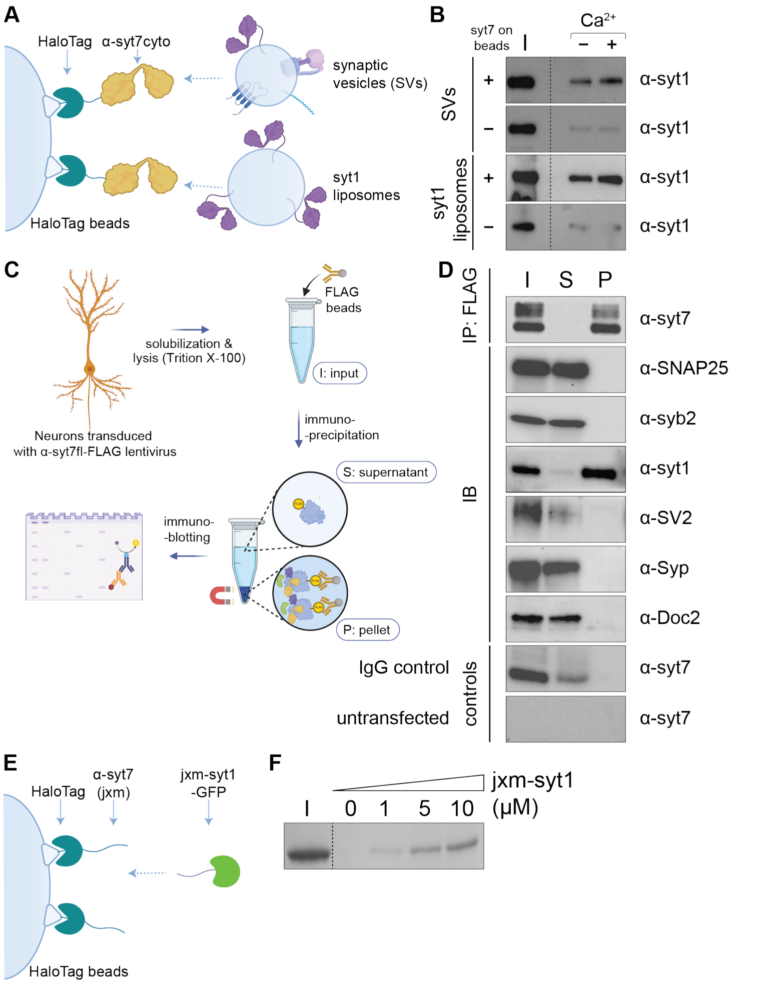

**Supplementary Figure 19. Direct interaction between the jxm linkers of α-syt7 and syt1, and α-syt7 directly binds purified synaptic vesicles (SV).**

(**A**) Schematic of the immunoprecipitation (IP) procedure showing neurons transduced with α-syt7-fl-FLAG lentivirus. The lysates (input) were incubated with anti-FLAG-conjugated magnetic Dynabeads to selectively pull-down α-syt7FLAG and its binding partners (pellet). IP'ed α-syt7FLAG, and potential binding partners were detected via Western blot analysis. The IP supernatant was included to assay for depletion of any bound species. (**B**) Blots showing successful pull-down of α-syt7FLAG. Only syt1 co-IP'ed; SNAP-25, syb2, SV2, Syp, and Doc2 failed to bind, suggesting a specific interaction between α-syt7FLAG and syt1. Two controls were used: IgG-conjugated Dynabeads and untransfected neurons; α-syt7-FLAG did not IP in either case. (**C,D**) Schematic and representative immunoblots from a HaloTag pull-down assay showing that α-syt7cyto directly binds to both purified SVs from mouse brain, as well as syt1-fl reconstituted into liposomes, in a weakly $Ca^{2+}$-dependent manner (SVs: 1.34 ± 0.23, syt1 liposomes: 1.46 ± 0.15). Empty beads showed minimal binding. (**E,F**) Schematic and representative gels from a HaloTag pull-down assay showing that direct binding occurs between the jxm linkers of syt7 and syt1 in a concentration-dependent manner. N=3 with three independent repeats for IP, and HaloTag pull-down assays, data are represented as mean ± SEM.

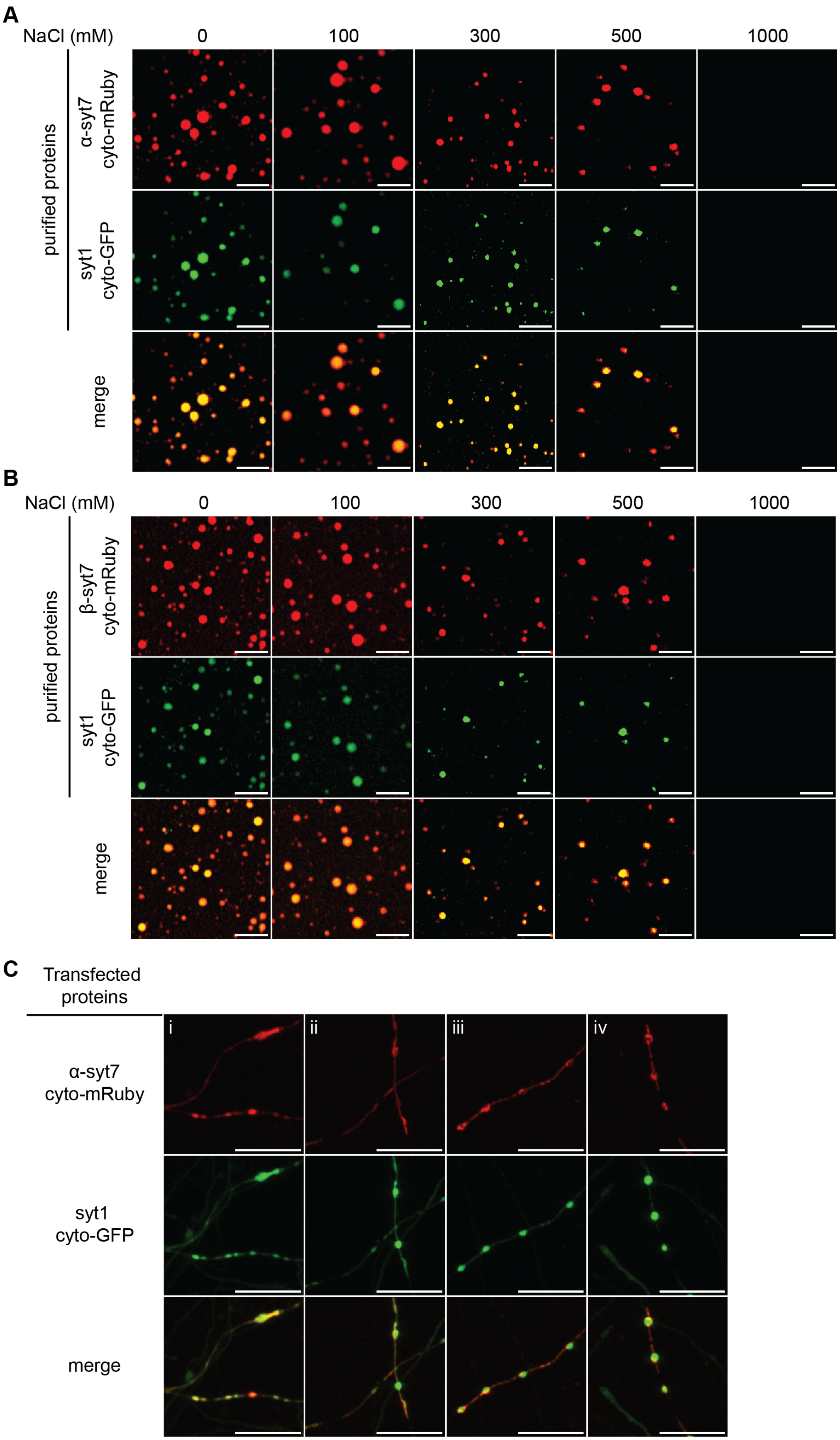
A
NaCl (mM)
0
100
300
500
1000
purified proteins
α-syt7
cyto-mRuby
syt1
cyto-GFP
merge
B
NaCl (mM)
0
100
300
500
1000
purified proteins
β-syt7
cyto-mRuby
syt1
cyto-GFP
merge
C
Transfected
proteins
i
ii
iii
iv
α-syt7
cyto-mRuby
syt1
cyto-GFP
merge

**Supplementary Figure 20. Syt7 and syt1 droplets coalesce *in vitro* and in hippocampal neurons.**

(**A**) Representative confocal image showing α-syt7cyto and syt1cyto droplets coalescing in the 0 mM NaCl condition. With increasing ionic strength, the coalescent α-syt7cyto and syt1cyto droplets dissolve, suggesting ionic interactions within droplets. Scale bar, 5 μm. (**B**) Same as (A), but for β-syt7cyto. Scale bar, 5 μm. (**C**) Representative images (i-iv) of rat hippocampal neurons transfected with α-syt7cyto-mRuby and syt1cyto-GFP illustrating that these droplets coalesce in neurites. Scale bar, 15 μm. N=3 with three independent repeats. The buffer used in (A,B) was 25 mM Tris-HCl (pH 7.4), indicated NaCl, and 3% PEG 8000.

**Supplementary Table 1.**

Sequences of alternative splice isoforms of syt7

| Syt7 alternative splice isoforms | Juxtamembrane linker protein sequence | # of residues |
|---|---|---|
| α-syt7 | **C**HW**C**QRKLGKRYKNSLETVGTPDSGRGRGEKKAIKL PAGGKAVNTAPVPGQTPHDESDRRTETRSSVSDLVNS LTSEMLMLSPGSEEDEAHEGCSRENL | 99 |
| β-syt7 | **C**HW**C**QRKLGKRYKNSLETVGTPDSGRGRGEKKAIND LDRDFWNNNESTVQQKWSSYPPKEFILNISPYAPYGD PRLSLKLPAGGKAVNTAPVPGQTPHDESDRRTETRSS VSDLVNSLTSEMLMLSPGSEEDEAHEGCSRENL | 143 |
| γ-syt7 | **C**HW**C**QRKLGKRYKNSLETVGTPDSGRGRGEKKAINF EDSTLSTATTLESIPSSAGEPKCQRPRTLMRQQSLQQ PLSQNQRGRQPSQPTTSQSLGQLQAHAASAPGSNP RAYGRGQARQGTSAGSKYRAAGGRSRSNPGSWDH VVGQIRNRGLDMKSFLEGRMVVLSLVLGLSEQDDFAN IPDLQNPGTQQNQNAQGDKRLPAGGKAVNTAPVPGQ TPHDESDRRTETRSSVSDLVNSLTSEMLMLSPGSEED EAHEGCSRENL | 263 |

Colored sequences of β- and γ-syt7 juxtamembrane linkers indicate changes in amino acid composition with respect to α-syt7. C, in bold, indicates cysteine residues which undergo palmitoylation.

**Supplementary Table 2.**

Kinetics of FRAP recovery of each of the three alternative splice variants of syt7 at the plasma membrane in HEK293T cells

| | Plasma membrane | |
|---|---|---|
| | Control | 1,6-HD treated |
| α-syt7 | 5.43 (3.98, 6.88) | 1.12 (-0.136, 2.38) |
| β-syt7 | 11.6 (6.37, 16.8) | 5.48 (2.20, 8.76) |
| γ-syt7 | 2.75 (0.479, 5.01) | 5.06 (0.630, 9.49) |

Calculated $t_{1/2}$ (s) values from fitting FRAP recovery curves with a hyperbolic function for each of the three alternative splice variants of syt7 in HEK293T cells, under control and 1,6-HD treated conditions. Data are represented as mean and 95% confidence interval (CI).

**Supplementary Table 3.**

Kinetics of FRAP recovery of each of the three alternative splice variants of syt7 at interbouton and bouton in rat hippocampal neurons

| | Interbouton | |
|---|---|---|
| | Control | 1,6-HD treated |
| α-syt7 | 15.3 (12.2, 18.4) | 8.63 (1.70, 15.6) |

| | | |
|---|---|---|
| β-syt7 | 21.8 (15.0, 28.6) | 5.98 (0.497, 11.5) |
| γ-syt7 | 8.3 (1.01, 15.6) | 13.6 (-1.07, 28.3) |

| | Bouton | |
|---|---|---|
| | Control | 1,6-HD treated |
| α-syt7 | 6.88 (0.40, 13.4) | 10.1 (2.97, 17.1) |
| β-syt7 | 4.42 (0.0121, 8.84) | 4.37 (0.0165, 8.72) |
| γ-syt7 | 115 (40.9, 190) | 44.2 (14.8, 73.6) |

Calculated $t_{1/2}$ (s) values from fitting FRAP recovery curves with a hyperbolic function for each of the three alternative splice variants of syt7 at interbouton and bouton in rat hippocampal neurons, under control and 1,6-HD-treated conditions. Data are represented as mean and 95% confidence interval (CI).

**Supplementary Table 4.**

Kinetics of decay of PPF tuning curves for the three syt7 splice variants

| Time Constant | WT | α-syt7 | β-syt7 | γ-syt7 |
|---|---|---|---|---|
| τ1 (ms) | 129 ± 19.5 | 143 ± 17.8 | 212 ± 12.3 | 50 (fixed; 56%) |
| τ2 (ms) | - | - | - | 1280 ± 315 (44%) |
| $R^2$ | 0.9954 | 0.9937 | 0.9988 | 0.9865 |

Calculated decay time constants (τ) and the coefficient of determination values ($R^2$) of PPF tuning curves for WT and syt7KO neurons expressing each of the three alternative splice variants of syt7. Data are represented as mean ± SEM. Note that WT, α-, and β-syt7 were fitted to a single exponential function, while γ-syt7 was fitted to a double exponential, with a fixed τ1. The amplitudes of the fast and slow components of decay for γ-syt7 are indicated with their respective τs. We restricted the faster-decay component of γ-syt7 to the lowest time point in our dataset; hence, the currently reported faster-decay constant is an underestimate.

**Supplementary Table 5.**

Mean and amplitude for nearest neighbor distance calculations

| Nearest neighbor distance (NND) | Gaussian peak | Mean ± SEM (nm) | Amplitude ± SEM |
|---|---|---|---|
| syt7 to syt1 | peak 1 | 22.8 ± 0.42 | 82.5 ± 4.22 |
| | peak 2 | 149 ± 3.13 | 29.8 ± 1.89 |
| syt1 to syt7 | peak 1 | 22.4 ± 1.39 | 33.4 ± 4.02 |
| | peak 2 | 160 ± 2.83 | 29.6 ± 2.51 |

Mean and amplitude of two peaks fitted with Gaussian functions for syt7 to syt1 and syt1 to syt7 nearest neighbor distance (NND) frequency distribution graphs in Fig. 6E,F, respectively. Data are represented as mean ± SEM.

**Supporting Material and Methods**

**DNA constructs**

Intensity-based glutamate-sensing fluorescent reporter (iGluSnFR) S72A was a gift from Looger L. (Janelia Farm, Ashburn, VA; AddGene ID# 106176)[4], but was modified with CAMKIIα promoter as previously described[1]. α-syt7 cDNA was a gift from Fukuda M. (Tohoku Neuroscience Global COE, Sendai, Japan)[5]. β- and γ-syt7 sequences were ordered as geneblocks from Integrated DNA Technologies; all three splice variants α, β, and γ-syt7-full length (fl) were assembled using PCR splicing with overlap extension and subcloned into transfer plasmid FUGW with human synapsin (hSyn) promoter and WPRE (Woodchuck hepatitis virus posttranscriptional regulatory element) 3' untranslated region (UTR) element. FUGW was a gift from Baltimore D. (California Institute of Technology, Pasadena, CA; AddGene plasmid #14883)[6]. pLenti-hSyn-CRE-WPRE plasmid was used for CRE expression (a gift from Fan Wang; Duke University Medical Center, Durham, NC)[7]. Constructs encoding the complete cytoplasmic domains of the alternative splice variants of syt7 were modified to add an N-terminal 6XHis tag, and a C-terminal fluorescent tag, mRuby3 (referred to as mRuby in the text). These fusion constructs were subcloned into the pET28(a)+ plasmid for bacterial expression. Rat syt1 cDNA was provided by T. C. Sudhof (Stanford University, Stanford, CA)[8]; the D374 mutation was corrected by replacement with a glycine. Constructs encoding SUMO-syt1-fl, syt1 cytoplasmic domain fused with GFP, and syt1 juxtamembrane linker fused with GFP were subcloned into pET28(a)+ plasmid for bacterial expression.

**Recombinant protein expression**

Recombinant proteins were expressed as described previously[9]. Briefly, all recombinant proteins were expressed in *E. coli* BL21(DE3) (NEB, C2527H) cells at 37°C until an OD of 0.6 was achieved. Bacteria were then induced with 500 µM isopropyl β-D-1-thiogalactopyranoside (IPTG) (GoldBio, I2481C) followed by overnight growth at 18°C. Cells were harvested and lysed using sonication, followed by solubilization with 1% Triton TX-100 (Thermo Fisher Scientific, A16046) for 2 h. A two-step protein purification, involving affinity and size-exclusion chromatography, was performed using Cobalt TALON affinity resin (Takara, 635653) and a Superdex 200 Increase 10/300 GL column (Cytiva, 28990944) on a fast protein liquid chromatography instrument (FPLC; Akta). Proteins were eluted in 25 mM Tris pH 7.4 buffer with 500 mM NaCl. To minimize proteolytic activity, a protease inhibitor cocktail (PIC) (Roche, 046693132001) was used during the lysis steps. High salt washes (0.5-1 M NaCl) were performed to remove any contaminants. For syt1-fl protein purification, similar steps plus 0.9% CHAPS detergent (3-((3-cholamidopropyl) dimethylammonio)-1-propanesulfonate; Fisher Scientific, 50-223-7023) were added during lysis and maintained throughout the purification steps. Syt1-fl was cleaved from the beads with SUMO protease treatment. Finally, purified proteins were subjected to SDS-PAGE, and protein concentration was determined using bovine serum albumin (BSA) (Jackson ImmunoResearch, 001-000-162) as a standard.

**Self-association assay**

*In vitro* self-association assays were conducted as described previously[9]. In short, mRuby-fused proteins (Fig. 1D, Supplementary Fig. 1A-F) were assessed for self-association in 25 mM Tris pH 7.4, 100 mM NaCl buffer conditions, at varying [protein] and [PEG 8000] (Sigma, P2139). In Fig. 1J,K, [NaCl], and [$Ca^{2+}$] were varied. Protein droplets and aggregates were imaged using a ZeissAxioVert.AX10 and Zeiss 880 Airyscan LSM microscope with a 63X/1.4 NA oil objective at room temperature (RT). Ten µl of each sample was placed on an 18 mm coverslip (Warner Instruments, 64-0734, CS-18R17), and the settled droplets and aggregates were imaged. Images were analyzed using the threshold and analyze functions in Fiji. Data were plotted using GraphPad Prism.

**HEK293T and neuronal cell culture**

HEK293T cells (ATCC, CRL-11268) were maintained in Dulbecco's Modified Eagle Medium (DMEM) with high glucose (Gibco, 11965092), supplemented with 10% fetal bovine serum (FBS; R&D Systems, S11550H) and penicillin-streptomycin (Thermo Fisher Scientific, MT-30-001 CI), as described previously[9]. Hippocampal neurons were isolated from pre-natal Sprague-Dawley rats (Envigo) on E18. HEK293T cells and rat neurons were plated on 18 mm coverslips (Warner Instruments; 64–0734 (CS-18R17)) that had been coated with poly-D-lysine (Thermo Fisher Scientific, ICN10269491) for 1 h at RT, at a density of 100K (HEK293T cells) or 125K (rat hippocampal neurons) per coverslip, in supplemented DMEM.

Mouse hippocampal neurons from *syt7$^{fl/fl}$* mice (#C57BL/6N-Syt7$^{em1(IMPC)H}$/H; repository ID: EM:14672; Mary Lyon Centre at MRC Harwell, which is the UK node of the European Mouse Mutant Archive (EMMA))[10–13] maintained as homozygotes, were isolated between P0-P1. Hippocampal tissue was dissected and maintained in chilled Hibernate-A media (BrainBits; HA). Post-dissection, neuronal tissue was incubated in 0.25% Trypsin (Corning; 25–053 CI) for 30 min at 37°C, and washed 2x with DMEM supplemented with 10% FBS, and Penicillin-Streptomycin. Tissue was triturated with a 1 ml pipette tip (1 mm diameter) until homogeneous. Cells were then counted with a Scepter 3.0 automated cell counter (Millipore Sigma; PHCC340) loaded with a 40 µm tip; gated to a cell diameter of 7-12 µm, and plated in DMEM media on 18 mm glass coverslips coated with poly-D-lysine and EHS laminin (Thermo Fisher Scientific; 23017015) at a density of 250K cells per coverslip. Both rat and mouse hippocampal neurons were incubated for 1 h at 37°C, 5% $CO_2$, after which DMEM media was aspirated and immediately replaced with neurobasal media-A (NBM-A, Thermo Fisher Scientific; 10888–022) supplemented with 2% B-27 (Thermo Fisher Scientific; 17504001) and 2 mM Glutamax (Gibco; 35050061). Following plating, neurons were fed with NBM-A plating media once a week.

**Transfection and transduction**

Lipofectamine-based transfection (Thermo Fisher Scientific, 15338-100) was carried out as previously described[9]. Lentivirus production and transduction were performed as previously described[1]. Lentivirus expressing CRE was added to neuronal cultures at 1 day in vitro (DIV), while lentivirus expressing iGluSnFR S72A and syt7 isoforms was added at 5-6 DIV. All viruses were titered by either fluorescence or Western blot prior to usage.

**Airyscan imaging and fluorescence recovery after photobleaching (FRAP) experiments**

HEK293T cells and cultured rat hippocampal neurons were imaged in standard ECF imaging solution at 37°C and 5% $CO_2$. Temperature, $CO_2$, and humidity were controlled using an Okolab incubation system (Okolab, Bold Line, Italy). FRAP was carried out on protein droplets/aggregates formed *in vitro*, and in HEK293T cells and rat hippocampal neurons using the photobleaching and time series modules of a Zeiss 880 Airyscan LSM microscope with a 63X/1.4 NA oil objective, using Fast Airyscan mode at RT (*in vitro* droplets) and 37°C (cell-based experiments). Briefly, we bleached circular regions of interest (0.8-2 µm in diameter) within protein droplets (2–2.5 µm in diameter) and aggregates using 405 and 488 laser lines at 25% and 100% laser power, respectively. Imaging was performed at 15 frames per minute. Samples were monitored for 20 s, 500 ms, and 400 s during pre-bleaching, bleaching, and recovery, respectively. To test for reversible dissolution of droplets, HEK293T cells and cultured rat hippocampal neurons were gently treated with 1,6-hexanediol (1,6-HD; Sigma, 88571) to a final concentration of 10% (v/v) for 10 min. Bleaching and imaging were performed in a manner similar to that described above. All images were processed with automatic Airyscan deconvolution. We normalized the fluorescence traces using the equation:

$$FRAP\ (t) = \frac{F.I._{bleach}\ (t) - F.I._{background}\ (t)}{F.I._{non-bleached}\ (t) - F.I._{background}\ (t)}$$

where, *F.I.* indicates fluorescence intensity. We performed FRAP experiments and averaged the *F.I.* data to obtain a single FRAP curve. Data were represented as mean ± SEM.

**Dynamic light scattering (DLS)**

DLS was carried out using a DynaPro Nanostar II Dynamic Light Scattering instrument (Waters Wyatt Technology) as described previously[9], using 10 µM protein buffered in 25 mM Tris pH 7.4, 100 mM NaCl, and 3% PEG 8000. Average diameter distributions were modeled using Rayleigh Spheres in the DYNAMICS v8 (Waters Wyatt Technology) software package. Samples were assayed in triplicate using three independent preparations, and the results were presented as mean ± SEM.

**Immunoblotting**

Immunoblotting was carried out as previously described with modifications[1]. Briefly, cell cultures were washed with 1x ice-cold PBS (Thermo Fisher scientific, 14190136) and lysed with 150 µl of lysis buffer (1x PBS, 2% SDS, 1% Triton X-100, 10 mM EDTA plus protease inhibitors: PMSF (Sigma-Aldrich, 11359061001), aprotinin (Sigma-Aldrich, A6103), leupeptin (Sigma-Aldrich, L9783), and pepstatin A (Sigma-Aldrich, P5318)). Lysed cell material was subjected to centrifugation at 20,000 x g for 30 min at 18°C, and the supernatant was collected. Lysates were then subjected to a bicinchoninic acid (BCA) assay (Thermo Fisher Scientific, 23227). After total protein quantification, a final concentration of 1x Laemmli Buffer (Bio-Rad, 1610747) plus 5% β-mercaptoethanol (BioRad, 1610710) was added to lysates, which were subsequently boiled at 100°C for 3 min. For protein detection, 5 µg of total protein was subjected to SDS-PAGE using 4-20% TGX-Stain Free gels (Bio-Rad, 5678094), and the gels were imaged on a Chemi-Doc MP system (Bio-Rad) following a 5 min activation time, and the resulting image was used as a total protein loading control. Protein gels were transferred to a PVDF membrane (EMD Millipore, IPFL00010) for 30 min per gel at constant current (240 mA), then blocked with 5% nonfat milk protein in Tris-buffered saline plus 1% Tween 20 (TBST) for 30 min.
PVDF membrane was incubated with primary antibody in 2.5% non-fat milk protein/TBST overnight. After three 10 min washes using TBST, the secondary antibody in 2.5% non-fat milk protein in TBST was incubated at RT for 1 h. After three 10 min washes with TBST, blots were incubated with a rabbit secondary antibody-HRP conjugate (BioRad, 1721019) for 1 h at RT. Blots were again washed thrice for a total of 30 min. Immunoblots were imaged using Luminata Forte Western HRP substrate (EMD Millipore; ELLUF0100) and a ChemiDoc MP Imaging System (Bio-Rad Laboratories). Bands were analyzed by densitometry, and contrast was linearly adjusted for publication using Fiji. A detailed list and sources of antibodies are included in the 'Antibody' section.

**Immunocytochemistry (ICC)**

Dissociated mouse hippocampal neuronal cultures were fixed with ice-cold 100% methanol, permeabilized with 0.2% saponin (Sigma Aldrich, 47036), blocked with 0.04% saponin, 10% goat serum (AbCam, ab7481), and 1% BSA in PBS, followed by immunostaining with primary antibody at 4°C overnight. Coverslips were washed with PBS three times and stained with secondary antibody in 0.1% BSA and 0.04% saponin in PBS for 1 h. Following three more PBS washes, the coverslips were mounted on microscope slides (Thermo Fisher Scientific, 22-178277), using ProLong Glass Antifade with Mountant with NucBlue Stain (Thermo Fisher Scientific, P36981), and imaged. A detailed list and sources of antibodies are included in the 'Antibody' section.

**iGluSnFR imaging and quantification**

iGluSnFR imaging and quantification were performed as previously described[1,2] using iGluSnFR S72A[4] with minor modifications. Briefly, experiments were performed on an

Olympus IX83 inverted microscope using an X-Cite 120 LED (Lumen Dynamics) and ORCA-Fusion CMOS camera (Hamamatsu Photonics). For high-frequency stimulation (HFS), neurons were depolarized with 50 action potentials (APs) at a frequency of 20 Hz (total of 2.5 s) by field stimulation, and 350 frames with 10 ms exposure at 2x2 binning were collected (total of 3.5 s). For paired-pulse stimulation, neurons were depolarized for two action potentials at a frequency of either 20Hz, 10Hz, 5Hz, 2Hz, or 1Hz, and 150 (20-2Hz) or 200 (1Hz) frames with 10ms exposure at 2x2 binning were collected (1.5-2s respectively). Extracellular imaging media was standard ECF (140 NaCl, 5 KCl, 2 CaCl2, 2 MgCl2, 5.5 glucose, 20 HEPES, pH 7.4, in mM) with an osmolarity of 300-320 mOsm. Imaging media was supplemented with 50 µM D-AP5 (Abcam, ab120003), 20 µM CNQX (Abcam; ab120044), and 100 µM Picrotoxin (Tocris; 1128) to block recurrent activity. All experiments were carried out at 33-34°C, with neurons between 14-15 DIV. Temperature and humidity were controlled by a Tokai incubation controller and chamber (Tokai Hit, PPZI3). A custom ImageJ plugin was used to identify iGluSnFR regions of interest (ROIs)[1,2]. Results from the plugin were imported to AxographX1.8.0 (Axograph Scientific), where traces were normalized to the initial 500 ms pre-stimulus baseline and corrected with a two-tailed background subtraction. Glutamate release signals were defined as fluorescence that was >4x the standard deviation (SD) of the noise. Due to the suppression of glutamate release in neurons expressing γ-syt7, it was difficult to analyze recordings at low frequencies. To adjust for this, the signal threshold was increased, and only ROIs with a signal >4x SD were used for further analysis. The synchronous fraction (within 10 ms of an AP) of the iGluSnFR signal was measured from multiple ROIs in each field of view (FOV). For spatial analysis, ROIs corresponding to the peaks from glutamate release signals from the 1$^{st}$ and 2$^{nd}$ APs were identified, color-coded, and superimposed on the average pre-stimulation FOV. PPF decay curves were fitted with either a single (WT, α-, and β-syt7) or double exponential function (γ-syt7), and are reported in Supplementary Table 4. Note that the faster-decay component of γ-syt7 is restricted to the lowest time point in our dataset. For the decay constant analyses in Supplementary Table 4, the Python analysis code was drafted using Claude Code (Anthropic, v1.15200.0 250bae, Sonnet 4.6). All scripts were reviewed, modified, and validated by the authors before use. The final code is available in the GitHub repository.

**STED super-resolution imaging and analysis**

Confocal and STED super-resolution imaging was performed with an Abberior MIRAVA Polyscope with the MATRIX module on an Olympus IX83 stand (Evident). The following hippocampal neuronal cultures were processed: WT, syt7KO, and syt7KO transduced with α-, β-, or γ-syt7-FLAG, with expression levels similar to those in Fig. 9B. Fixation, permeabilization, blocking, washing, and mounting were performed as described in ICC. The following primary and secondary antibodies were used: anti-FLAG (CST, D6W5B), anti-syt1 (DSHB, mAb48), and anti-bassoon (SYSY, 141005); anti-rabbit IgG-coupled

Abberior STAR RED (Abberior, STRED-1002), anti-mouse IgG-coupled Alexa 594 (Thermo Fisher Scientific, A11005), and anti-guineapig IgG-coupled Alexa 488 (Thermo Fisher Scientific, A110073). 3D STED images were acquired with an oil 60X objective NA:1.42 (Evident, UPLXAPO60XO), 488, 561, and 640 nm excitation laser lines, 775 nm STED depletion laser (~20%), with a pixel size of 50 nm. Images were analyzed using Abberior Lightbox 2025 and Fiji. Synapses with a round bassoon (ellipticity ≥ 0.8) and overlapping syt1 or syt7 signal were classified as en face synapses. Manual thresholding was applied to filter bright signals. Syt1 and syt7 clusters were identified (radii ≥ 3 pixels) and counted in bassoon-positive areas.

**MINFLUX imaging and image analysis**

For sample preparation, mouse hippocampal neurons expressing α-syt7-FLAG were fixed and permeabilized as described in ICC. Syt1, FLAG-tag, and bassoon were labeled with primary (see Antibodies section) and secondary antibodies (FX640, FX680, and Alexa 488). We note a ~20 nm linkage error due to primary and secondary antibody labeling.

To stabilize the sample during measurements, gold nanoparticles (BBI Solutions, EM.GC150) were used as fiducial markers[14,15]. These nanoparticles were applied to the samples, and any unbound particles were washed away with PBS. For imaging, a buffer containing glucose oxidase, called GLOX buffer, was used (50 mM Tris-HCl pH 8.0, 10 mM NaCl, 10% (w/v) glucose, 64 µg/ml catalase, 0.4 mg/ml glucose oxidase, and 25-50 mM mercaptoethylamine (MEA)). After mounting, samples were sealed with twinsil (picodent). Images were collected using Abberior Instruments Imspector software (version 16) with MINFLUX drivers, using a 100X magnification NA 1.4 oil objective lens. Stabilization was achieved using a linearly polarized 980 nm laser (Thorlabs Inc., LP980-SF15). After selecting a FOV, 75 × 75 µm confocal images were acquired. For activation of single fluorophores, a 405 nm wavelength laser (HÜBNER Photonics, Cobolt MLD 405 nm 50 mW) whose intensity was attenuated into the nW region by a neutral-density filter. Photons emitted from the sample were counted using two avalanche photodiodes.

Data acquired from Abberior Imspector software were saved as .MSR and exported into the .NPY format. These raw data were processed to optimize EFO, CFR, and len_min with values ranging from 100-300 kHz, 0-0.9, and >3, respectively (Note: the definitions of these parameters are defined below). For spectral unmixing of the two fluorophores, DCR was fitted with two Gaussian functions to separate FX640 and FX680. The mismatch in the refractive index between the #1.5 coverglass and the aqueous imaging buffer (GLOX buffer) distorts the position measurement along the Z-coordinate. This effect makes the focal point appear shallower than it actually is. To correct for this distortion in standard samples, a correction factor of 0.7 was applied to the measured Z-position. To find the nearest neighbor distance (NND), the distances between color-coded TIDs were

calculated and sorted. To obtain confocal images of bassoon, .MSR files obtained through FIJI, and the centroid of all bassoon puncta was calculated (thresholding via the Otsu method, radius range in pixels 3-20, overlap threshold 0.75, and eccentricity threshold 0.9375)[16]. Relative positions of every TID in the 640 and 680 DCR split were updated based on the bassoon centroids, and all the localizations were merged. Clustering was performed using density-based spatial clustering of applications with noise (DBSCAN). This algorithm groups together points that are closely packed, marking outliers as points that lie alone in low-density regions. It works by identifying "core points" with a minimum number of neighbors (5-8 in this case) within a certain radius (22.5 nm in this case), and then expanding clusters outward from these points. Any point that is not reachable by this expansion is labeled as noise, allowing the algorithm to find arbitrarily shaped clusters and effectively handle outliers without a predefined number of clusters. After clustering, all the clusters within 300 nm of the centroid were plotted in 3D and 2D projections in XY, XZ, and YZ. All analysis was done using Python.
The relevant definitions for this section of Methods are as follows:

- Effective frequency at offset (EFO): effective emission frequency (in Hz) measured at offset pattern positions.
- Center frequency ratio (CFR): the emission frequency measured at the center position of the pattern (EFC) is divided by the emission frequency measured at the offset pattern positions (EFO). CFR = EFC / EFO.
- Detection channel ratio (DCR): The counts measured on ‘Gate Channel 1’ is divided by the counts measured on ‘Gate Channel 1’ and ‘Gate Channel 2’:
- Trace ID (TID) is trace ID. Consecutive localizations from a single event share the same trace ID.
- Len_min: minimum number of localizations per tid.

**Immunoprecipitation (IP)**
Magnetic Dynabeads M270 Epoxy (Thermo Fisher Scientific, 14302D) were covalently linked to the FLAG antibody (Rabbit FLAG M2 antibody, Sigma) as follows: 10 mg of beads were mixed with 250 µg of antibody in 400 µl of borate buffer (100 mm sodium borate, pH 8.5) plus an additional 200 µl of 3 M ammonium sulfate overnight at 37°C. Purified control rabbit IgG was also conjugated to Dynabeads as a negative control. After coupling, the beads were subjected to a series of six rigorous 1 ml washes with alternating pH buffers (500 mM NaCl, 50 mM ammonium acetate, pH 4.5, and 500 mM NaCl and 50 mM Tris-HCl, pH 8.0) using a magnetic stand to remove unbound antibody. Finally, the prepared antibody-coated beads were washed and resuspended at 30 mg/ml in 150 mM KCl and 50 mM Tris-HCl, pH 8.0, and stored at 4°C until needed.
Rat hippocampal neurons expressing α-syt7-fl-FLAG were lysed using lysis buffer (25 mM Tris-HCl, pH 7.4, 100 mM NaCl, 1 mM EGTA, 5% glycerol, 1% TritonX-100 along with a protease inhibitor cocktail (1 tablet/ 10 ml; Sigma Aldrich 11836170001). After

solubilizing on ice for 20 min with intermittent agitation, large insoluble cell debris was removed by centrifugation (4 min, 13,000 × *g,* 4°C). Supernatant was collected (Input fraction) and incubated with FLAG-Dynabeads on ice, with rotation, for 2 h. Beads were collected using a magnetic stand, and the supernatant fraction was stored. Beads were washed thrice with the lysis buffer, followed by elution with 4x Laemmli sample buffer (Bio-Rad, 1610747) with β-mercaptoethanol (Bio-Rad, 1610710). All samples were heated at 75°C for 10 min and subjected to SDS-PAGE and immunoblot analysis.

**Synaptic vesicle (SV) purification**

SV purification was performed as described previously[17], again using Magnetic Dynabeads M270 Epoxy (Thermo Fisher Scientific, 14302D) conjugated to the Rho1D4 antibody (purchased from the University of British Columbia (https://ubc.flintbox.com)) as described in the IP section.

SVs were isolated from WT mice generated from crossing synaptophysin[18] and synaptogyrin[19] breeder colonies at 14 - 20 days of age and of both sexes. Mice brains were isolated and homogenized in 125 mM KCl, 20 mM potassium phosphate, 5 mM EGTA, and protease inhibitors (cOmplete Mini EDTA-free, 1 tablet/10 ml; Sigma Aldrich 11836170001), pH 7.3) to create a crude lysate; all materials throughout the isolation procedure were maintained at 4°C. This lysate was centrifuged (20 min, 35,000 × *g*, 1°C) to remove large debris. Antibody-coated Dynabeads were incubated with 1.9 ml of the brain supernatant and incubated for 25 min with rotation on ice. The beads were collected using a magnetic stand, and the supernatant was discarded. Beads were then washed four times with an ice-cold wash buffer (145 mM KCl, 10 mM potassium phosphate pH 7.3), and SVs were eluted by incubating with 50 µl of 1 mM 1D4 peptide (Cube Biotech) for 30 min on ice in a buffer containing 100 mM sodium borate pH 8.5.

**Proteoliposomes and GUV preparation**

Syt1-fl was mixed with lipids (1 mM DOPC; Avanti Research 850375) on ice in 25 mM HEPES pH 7.4, 100 mM KCl, and 0.9% CHAPS (Fisher Scientific, 50-223-7023). Total volume was adjusted to bring the detergent concentration below the critical micelle concentration. Detergent was removed by overnight dialysis at 4 °C. Proteoliposomes were then isolated by flotation of the vesicles using an Accudenz (Accurate Chemical, AN7050/BLK) step gradient.

To produce giant unilamellar vesicles (GUVs), a thin film (~15 µl) of phospholipids (DOPC, DGS-NTA, and Atto647N DOPE in the ratio 93.5:5:1.5, 1 mM total; Avanti Research 850375, 790528; Sigma 42247) was deposited onto indium tin oxide (ITO) coated glass slides. Using Nanion Vesicle Prep Pro, the slides formed a chamber sealed by o-rings, which was filled with a 200 mM sucrose and 1 mM HEPES buffer solution. The electroformation process was initiated by applying a 10 Hz AC voltage of 3 V for 2 h at 37°C, which promotes hydration and self-assembly of the lipid film into vesicles. The resulting GUV solution was then collected, washed in an iso-osmolar buffer to remove residual sucrose and unincorporated substances, and filtered to isolate vesicles larger than 3 µm. Atto647N-labeled GUVs were incubated with α-syt7cyto-mRuby in 25 mM Tris pH 7.4, 100 mM NaCl, and 3% PEG 8000; α-syt7cyto droplets on GUV surface were imaged in a Bioinert µ-dish (Ibidi, 81150) using confocal microscopy. All consumables were brought from Nanion Technologies GmbH.

**HaloTag-pull down assay**

HaloLink resin was washed extensively before protein attachment, and blocked with BSA to prevent non-specific binding. Purified Halo-tagged proteins (~1.2 mg, 'bait' proteins) were immobilized on HaloLink resin (750 µl bed volume) for 1 h, at 25 °C, followed by 3x buffer wash. Buffer was 25 mM HEPES pH 7.4, 100 mM KCl, and indicated 0.2 mM EGTA or 1.2 mM $Ca^{2+}$. Complete binding was confirmed by SDS-PAGE analysis of the supernatant. Prey proteins at indicated concentrations were incubated with 'bait'-protein-tagged HaloLink bead slurry, such that the final concentration of the bait protein was 7 µM. This mixture was incubated for 1 h with rotation at 25°C and supernatants obtained by centrifugation (3500 × *g*, 10 min) were used for SDS-PAGE and Western blotting.

**Hydropathicity analysis**

The hydropathicity scores for the juxtamembrane linkers of α-, β-, and γ-syt7 (Supplementary Table 1) were calculated using: https://web.expasy.org/protscale/. These scores are based on Kyte and Doolittle scoring index. A hydropathicity score of 0.25 was used as a cut-off; a high hydropathicity index indicates high hydrophobicity.

**Antibodies**

Primary Antibodies:

| Antibody | Source | Identifier | Concentration |
|---|---|---|---|
| Anti-syt7<br>(Rabbit pAb) | SYSY | 105-173; RRID: AB_887838 | WB: 1:3000 |
| Anti-syt7<br>(Rabbit mAb) | AbCam | ab311843 | WB: 1:2000 |
| Anti-FLAG<br>(Rabbit mAb) | Cell Signaling Technology | D6W5B, 14793; RRID: AB_2572291 | WB: 1:1000<br>ICC: 1:500<br>IP: 1:40<br>MINFLUX: 1:2500<br>STED: 1:400 |
| Anti-Rho1D4<br>(Mouse mAb) | University of British Columbia | RHO 1D4 | IP: 1:40 |
| Anti-SNAP25<br>(Rabbit mAb) | AbCam | EPR3275; RRID: AB_10887757 | WB: 1:1000 |
| Anti-syb2<br>(Mouse mAb) | SYSY | 104 211; RRID: AB_2619758 | WB: 1:1000 |

| | | | |
|---|---|---|---|
| Anti-syntaxin (Mouse mAb) | SYSY | 110 011; RRID: AB_887844 | WB: 1:1000 |
| Anti-SV2 (Mouse mAb) | Developmental Studies Hybridoma Bank (DSHB) | SV2; RRID: AB_2315387 | WB: 1:500 |
| Anti-VGlut1 (Mousse mAb) | NeuroMab | VGlut1; RRID: AB_2187693 | ICC: 1:500 |
| Anti-Pan-Neurofascin (Mousse mAb) | NeuroMab | Pan-Neurofascin (extracellular); RRID: AB_2187693 | Live imaging: 1:100 |
| Anti-syt1 (Mouse mAb) | DSHB | MAB48 (asv 48); RRID: AB_2199314 | WB: 1:1000<br>MINFLUX: 1:2000<br>STED: 1:300 |
| Anti-Doc2a/b (Rabbit pAb) | SYSY | 174 203; RRID: AB_11064600 | WB: 1:250 |
| Anti-Bassoon (Guinea pig pAb) | SYSY | 141 005; RRID: AB_2924946 | ICC: 1:250<br>STED: 1:300 |
| Anti-Syp (Guinea pig mAb) | SYSY | 101 308; RRID: AB_2924959 | WB: 1:500 |

Secondary Antibodies:

| Antibody | Source | Identifier | Concentration |
|---|---|---|---|
| Goat anti-Rabbit IgG HRP | Bio-Rad | 1706515; RRID: AB_11125142 | WB: 1:10,000 |
| Goat anti-Mouse IgG-HRP | Bio-Rad | 1706516; RRID: AB_2921252 | WB: 1:10,000 |
| Goat anti-Mouse IgG2α-Alexa Fluor 647 | Thermo Fisher Scientific | A21241; RRID: AB_2535810 | ICC: 1:500 |
| Goat anti-Mouse IgG-Alexa Fluor 594 | Thermo Fisher Scientific | A11005; RRID: AB_141372 | STED: 1:200 |
| Goat anti-Rabbit IgG-Alexa Fluor 546 | Thermo Fisher Scientific | A11035; RRID: AB_143051 | ICC: 1:500 |
| Goat anti-Guinea Pig IgG-Alexa Fluor 488 | Thermo Fisher Scientific | A11073; RRID: AB_2534117 | ICC: 1:500<br>STED: 1:200 |
| Goat anti-Rabbit IgG-STAR RED | Abberior | STRED-1002; RRID: AB_2833015 | STED: 1:200 |

| | | | |
|---|---|---|---|
| Goat anti-Guinea Pig IgG HRP | AbCam | Ab6908; RRID: AB_955425 | WB: 1:10,000 |
| Goat anti-Mouse IgG-Flux 640 | Abberior | FX 640 | MINFLUX: 1:2500 |
| Goat anti-Rabbit IgG-Flux 680 | Abberior | FX 680 | MINFLUX: 1:3000 |

**Data S1. (separate file)**
Raw data and statistical analysis information for Figures 1-6.

**Data S2. (separate file)**
Raw data and statistical analysis information for Supplementary Figures 1-20.
